# Translation of small ORFs in non-canonical RNAs provides resilience to heat shock stress

**DOI:** 10.64898/2026.11.19.767816

**Authors:** Casimiro Baena-Angulo, Rosario Morrugares, Ana Isabel Platero, José Ignacio Pueyo, Ana Belén Hervás, Emile Magny, Juan-Pablo Couso

## Abstract

**Background:** Heat shock and other stress conditions severely challenge cell and organismal survival. Characterising and understanding the genome responses to heat shock is of increasing interest due to the accelerated impact of global warming. Under heat shock and other stress conditions, the translation of most canonical genes is reduced as part of a physiological response to improve cellular and organismal resilience. Heat shock proteins (HSPs) are among a minority of proteins whose expression increases during heat shock, working as chaperones and helping other proteins to maintain their tertiary structure.

**Results:** In contrast to canonical genes, whose translation is dramatically suppressed under heat shock conditions, we find that the translation of small Open Reading Frames (smORFs), especially those in so-called long-noncoding, (or long noncanonical) RNAs (lncRNAs), remains at similar levels, and in some cases is enhanced under heat shock in *Drosophila melanogaster*. lncRNA translation under heat stress entails a 3’ shift in the smORFs translated in a given lncRNA, a shift that may be favoured by non-canonical smORF translation mechanisms. We also characterise two smORFs with stress-enhanced translation, showing that their expression is up-regulated by the heat stress response pathway, and that their microproteins improve organism survival and resilience under heat shock.

**Conclusion:** smORF and lncRNA translation is more resilient to heat shock than that of standard genes, and this smORF translation can increase organismal resilience to heat shock, potentially representing a new component of the HSF stress-response gene regulation pathway. The stress-resilient lncRNA translation entails a 3’ shift in the position and translation levels of smORFs being translated, and while the full molecular mechanism of this shift is not known at the moment, we propose that it might take advantage of non-canonical aspects of polycistronic lncRNA translation.

## BACKGROUND

Small ORFs (smORFs) of less than 100 codons are at the interface between coding and non-coding genes, and their number (millions in the human genome) challenge our understanding of the coding content of our genomes [1, 2]. Yet, most of them remain uncharacterised and even non-annotated in most genomes, due to a mixture of historical reasons and technical difficulties. The short size of smORFs and their microproteins complicates genetics, molecular biology and biochemistry experiments; both their transcription and translation levels are usually low and present only in a cell- or stage-specific manner; their mutant phenotypes tend not to be overt morphological defects, but more subtle cellular and physiological failures [3, 4]; and finally, our collective ideas of what is a gene and an annotated metazoan genome, find it difficult to accept that we may have missed thousands of uncharacterised genes producing peptides as small as 11 amino acids (aa) [5].

We have previously proposed, using a combination of bioinformatic analysis and extrapolation from functional studies, that metazoan smORFs include several classes of sequences with different molecular characteristics and functions (Fig. 1A). smORFs range from randomly-arising yet inert sequences in DNA (intergenic ORFs), to regulators of gene expression — sometimes through their own translation (lncORFs), and finally to short coding sequences expressing functional peptides with propensity to act as regulators of canonical proteins (sCDSs). The characteristics of different smORF classes are compatible with representing steps in the evolution of novel coding genes from random DNA sequences [2, 6, 7]. In this emerging view, the function of smORFs as a whole would be to provide the genome with new elements for its continuing function and adaptation. However, the mechanisms whereby seemingly random non-coding sequences acquire coding function remain unclear. smORFs and novel genes must first achieve transcription, and then translation, and finally the translated peptides must prove of beneficial, adaptive value to be selected for and fixed [8–11]. We have provided a model [6] for the acquisition of coding functions from non-coding ones to explain the shift from long non-coding RNAs to RNAs translating functional smORF microproteins. In addition, it has also been proposed that stress, and other out-of-normal situations, provide opportunities for novel genes and microproteins to integrate into and improve fully-working genomes which are otherwise adapted to normal conditions [12–16].

**Figure 1.**
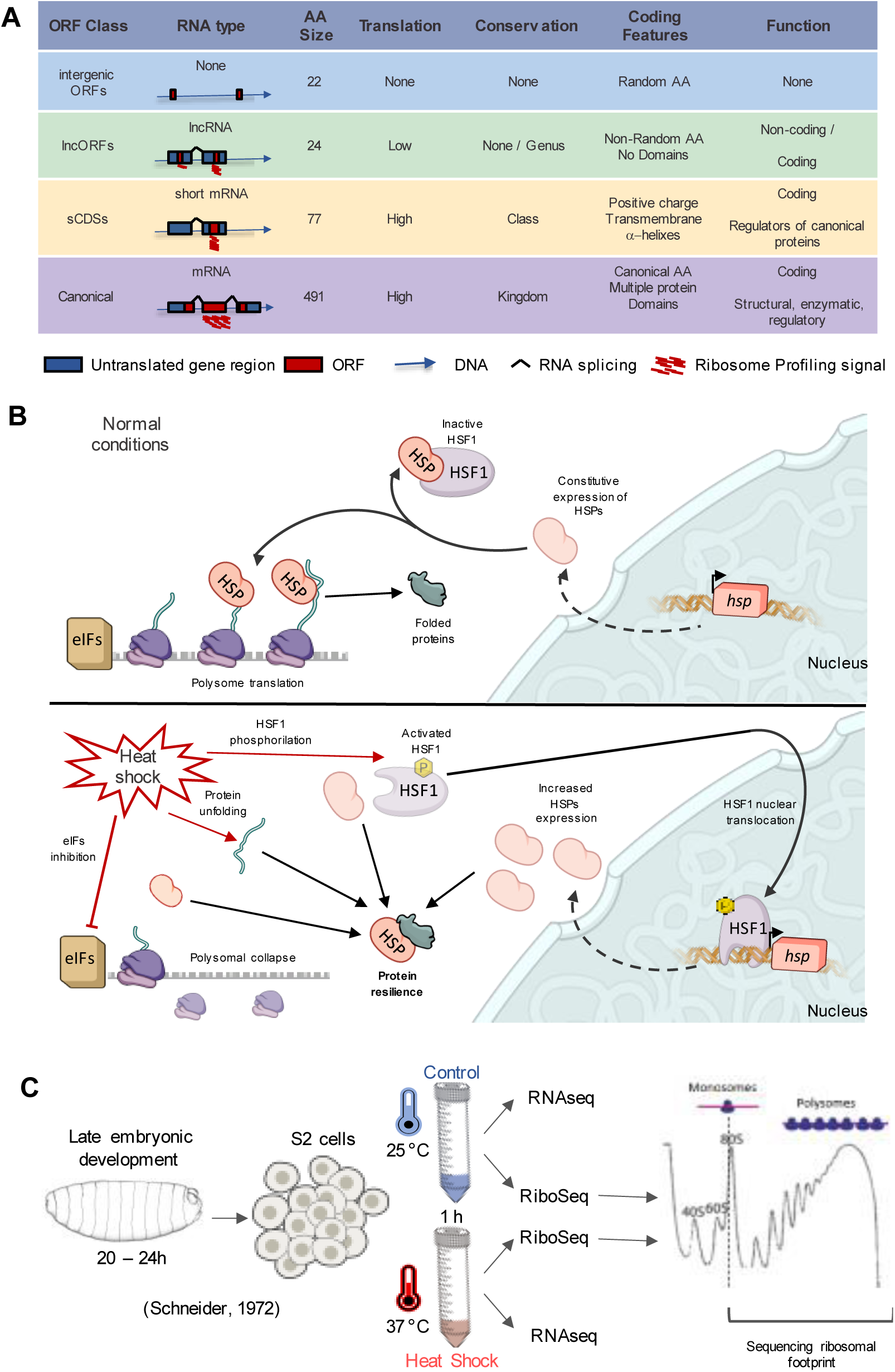
Overview of genomic and cellular context and experimental design. **A**. Types of ORFs relevant to this work and their properties, modified from Baena-Angulo et al., 2025 [6]. In animal genomes, most smORFs (ORFs < 100 codons) are placed randomly in untranscribed DNA regions (intergenic ORFs, red squares over blue arrow) or canonical ORFs (purple) those ORFs >101 codons which are present in their own mRNAs, that are processed and translated, and which are annotated as protein-coding, encoding conserved proteins (on average at the kingdom level) with a large variety of functions. Between these two extremes, we find two smORF classes; First, short coding sequences (sCDS; yellow), similar to canonical ORFs (annotated as coding, located in processed and putatively translated mRNAs) but < 101 codons in length. sCDSs do not arise from short spliced-isoforms of canonical genes, nor are they located in untranslated leaders of canonical mRNAs (as upstream ORFs and downstream ORFs do [87, 88]) but in their own, smaller mRNA. Microproteins derived from sCDSs are conserved at the class level and mostly involved in regulatory functions due to their small size. Second, we find lncORFs (green). They are smORFs located within exons of long RNAs annotated as non-coding (lncRNAs), They generally show no or low translation levels but can behave as polycistronic coding genes. Their putative coding functions are variable and mostly unknown, but they have non-random and non-canonical amino acid usage, and those translated are mostly only conserved at the genus level. **B.** Model of the cellular heat shock response through Heat Shock Factor 1 (HSF1) activation, transcriptional induction of Heat Shock Protein genes (HSP). Under normal conditions HSP-bound HSF1 remains inactive, while HSPs also ensure the proper folding of proteins as they are released from ribosomes involved in 5’cap-dependent translation. Upon heat shock stress, some eIFs (Eukaryotic Translation Initiation Factors) are inactivated, inducing the arrest of canonical 5’cap-translation and the collapse of polysomes along the first 200nt of the transcripts. Resilient monosomes involved in translation are now releasing misfolded proteins. In parallel, HSF1 becomes activate through its phosphorylation and uncoupling from HSPs. Active HSF1 (HSF1Ps) can translocate to the nucleus to trigger the expression of HSPs. Once in the cytoplasm, HSPs interact with the misfolded proteins, helping to restore their native tertiary structure. **C**. Experimental design of the heat shock experiment. S2 cellcultures were either subjected to 1 hour of heat stress (37 °C) or maintained at room temperature, and total RNA was immediately isolated for RNAseq, while monosomal, polysomal and whole cytoplasmic fractions were subjected to RiboSeq and sequenced by NGS.

Cells respond to different stresses, either intrinsic or environmental (including DNA damage, ER stress, redox, hypoxia, starvation and infection). In higher organisms, cellular stresses can trigger organ and organismal stresses, and likewise, responses to stress can involve different organs as the cells and the body try to recover homeostasis [17, 18]. Cellular and organismal stresses are important research fields for biology, biotechnology and medicine, as stresses and their responses are main elements of physiopathology. *Drosophila melanogaster* is a useful model to study cellular and organismal stresses: most of its organs offer simplified versions of human ones, while maintaining high degrees of functional homology, both at the physiological and gene regulation levels. *Drosophila*’s cell biology, molecular biology and genetics have been exploited for many years to contribute to general and applied knowledge. In particular, *Drosophila*’s response to heat shock (HS) is a classic model of cellular stress response with direct homologies from yeast to humans [18, 19].

Translation is an often-overlooked regulatory step that allows rapid modulation of gene expression under developmental and environmental changes [20–23], and in fact, the translational read-out measured by RiboSeq is at least as reproducible as the RNA expression data provided by RNAseq [24, 25], implying a similarly tight regulation. Translational regulation is critical during the cells’ response to stress, including heat shock [17, 26–29]. Heat shock reduces and reprograms translation of canonical genes in metazoans, as heat shock proteins (HSPs) involved in translation disengage from polysomes in order to act as chaperones, binding and protecting already formed proteins from heat-induced mis-conformations, at the expense of leaving nascent proteins unprotected (Fig. 1B). Interestingly these unprotected nascent proteins have been suggested to play a role in triggering widespread elongation pausing, through their own misfolding after reaching a length of around 65 aa [19, 27, 30]. Prolonged heat shock leads to severe reduction of translation (visualised by the collapse of polysomes, Additional file 1: Fig. S1). Simultaneously, transcription of HSPs is enhanced by the Heat Shock Factor (HSF), a transcription factor that is sequestered in the cytoplasm by HSPs under normal conditions but is released and translocated to the nucleus upon heat shock [19, 31–33]. Thus, the heat shock response appears to have two components: genome-wide translational repression, and the transcriptional upregulation of specific genes that enhance stress resilience.

Other types of stress utilise a similar pathway to down-regulate canonical gene expression and enhance expression of factors bringing about stress damage limitation and cellular and organismal resilience [34]. However, not all aspects of the cellular stress responses are understood; and while more research is needed, heat shock in particular is becoming a more pressing issue in the context of global warming-induced heat waves. A function for smORFs during cellular and organismal stress fits well with the proposed evolutionary roles of emerging genes [2, 7], with smORF translational plasticity [24], and is supported by functional studies at genomic scale in yeast [12, 14, 35] and plants [36]. Moreover, their small protein length, often smaller than 65 aa, may render them impervious to the elongation pausing that hinders the translation of canonical longer proteins during heat stress [27]. Here, we use RNAseq and Ribosomal Profiling (RiboSeq) to reveal the RNA and translational profiles under heat shock of canonical and smORF genes in the metazoan *Drosophila melanogaster*, and show that compared to canonical genes, smORF translation is resilient to heat stress. Further, our functional investigation of two of the highest-responsive smORFs, highlights a role for smORFs in mitigating heat shock stress. We propose that non-canonical aspects of smORF translation might favour microproteins production under heat shock, and we show that some of these smORF microproteins integrate into the canonical heat shock pathway in order to increase organismal resilience to stress. Thus, our results suggest that smORFs may represent a so far unexplored component of the heat shock response, which also has important implications for the functional understanding of smORF genes as a class, and for the function and evolution of novel genes.

## RESULTS

### Heat shock produces a drop in canonical translation

The fruit fly, *Drosophila melanogaster,* provides a genetically tractable model to study responses to heat shock and other stresses at the genomic level. Long-established and near-isogenic lines of wild type flies reduce genetic variability, but other types of variability affecting experimentally-induced stress responses, be it intrinsic (developmental and physiological states), or extrinsic (environmental and culture conditions) remain. These further sources of variability can be reduced by using cell culture lines derived from the isogenic wild type flies. Thus, we have studied the heat shock response in *Drosophila* S2 cell line cultures [37], which have been extensively used for decades for genetic and genomic studies, with results validated by *in vivo* studies [38–41]. We used RNAseq and RiboSeq to evaluate global changes in RNA expression and protein translation upon heat stress (1h, 37 °C) in S2 cell cultures. To increase experimental robustness, we obtained two independent biological replicas (Additional file 1: Fig. S2A), each composed of paired control and heat shocked samples (see Methods, Fig. 1C and Additional file 1: Fig. S2B), and analysed them similarly to our previously published works [24, 42].

Comparison of the two replicas reveals a very high consistency and reproducibility of RNAseq results for annotated protein-coding genes (Fig. 2A), showing Spearman correlations around 0.90 between control replicates for RPKM^RNA^ values (RNAseq reads per kilobase per million reads mapped). There was a 93.8% overlap between genes showing RNA expression (defined as having RPKM^RNA^ ≥1) in either control replica, with a total of around 12,500 annotated genes detected as transcribed under control conditions. The heat shocked RNAseq replicas show near identical reproducibility as the controls, with a near 0.91 correlation in RPKM^RNA^ values and a 92.5% overlap in canonical genes deemed transcribed, for a total of 12,795 transcribed canonical genes under heat shock (Fig. 2A-B; Additional file 1: Fig. S2B).

**Figure 2.**
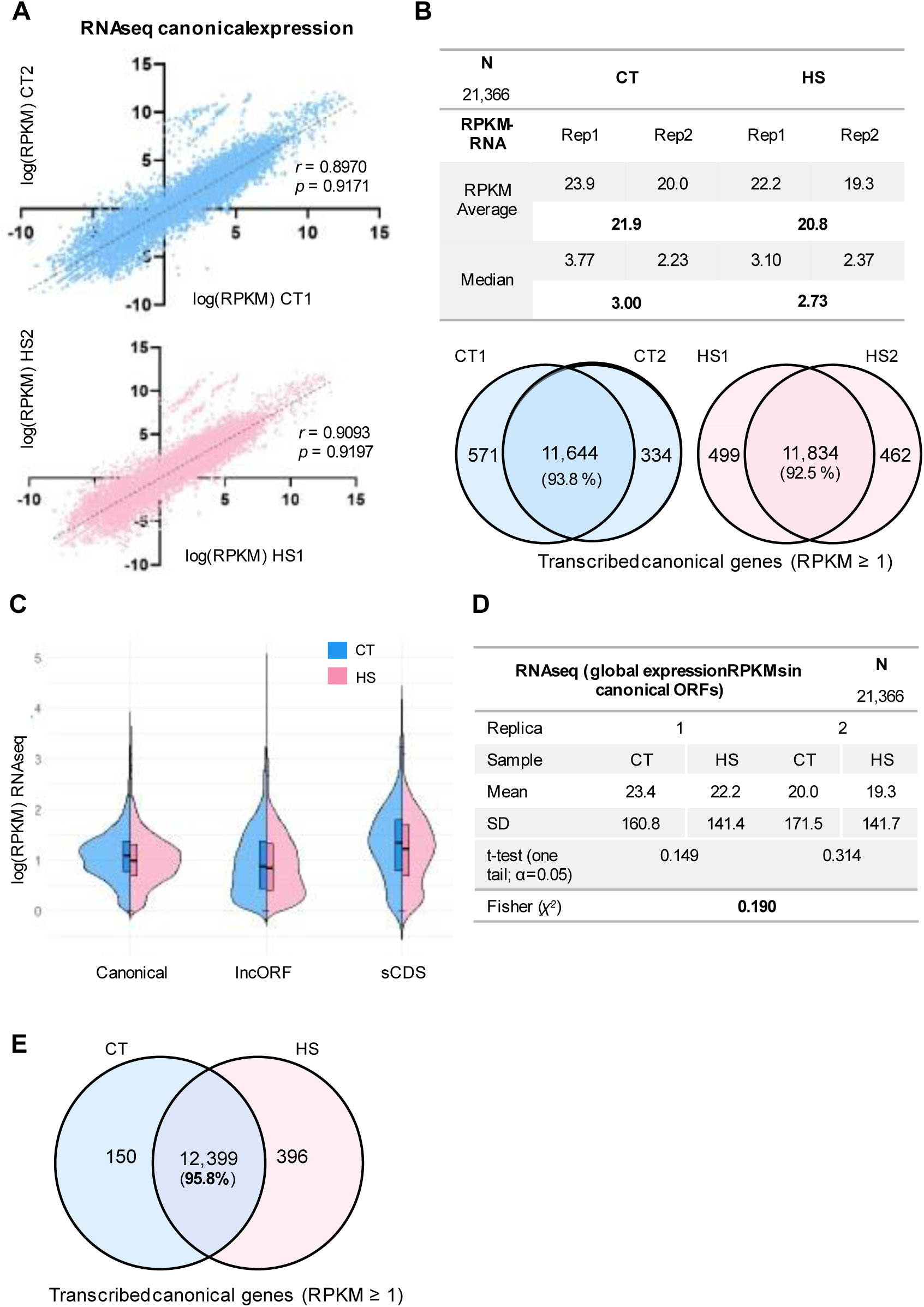
Resilient overall mRNA levels upon Heat Shock. **A.** High correlation of RNA expression (RNAseq) between replicates for all canonical genes in control (CT, blue) and heat shock conditions (HS, pink). (r: Spearman correlation; p: Pearson correlation). Dashed lines follow a slope x=y, indicating unchanged levels. **B.** Expression levels of canonical genes. The table shows average and median RNAseq expression (RPKMRNA) per replica and per condition (bold). Venn diagrams show the overlap of transcribed genes (those with RPKMRNA values ≥ 1) between replicas. **C.** Distribution of RNAseq expression levels across curated canonical ORFs, sCDSs and lncORFs between control and heat shock conditions, after filtering genes for their RNA expression values across replicates (see “New lncORFs annotation” in Methods section). RPKM values plotted for each ORF and condition are the averages between replicates. **D.** Summary of the expression levels of standard genes (see Additional file1: Fig.2B). The table shows mean RNAseq RPKM values for each condition in each replica, and the results of one-tailed t-tests showing significant variation between conditions for both replicas. Overall significance was assessed by Fisher’s method for combining p-values from two biological samples (see Methods). **E.** Overlap of all transcribed (RPKMRNA ≥ 1) canonical genes between control (CT) and heat shock (HS) conditions.

We observe no significant differences in overall average RNA expression of canonical ORFs between control and heat shock conditions (Fig. 2C-D), neither comparing a) the overlap of transcribed genes in control and heat shock conditions (Fig. 2E); or b) individual gene RPKM^RNA^ values in paired control-heat shock samples (*p*= 0.190 Fig. 2D; see Methods).

However, global averages without significant alterations do not exclude the transcriptional upregulation of some genes, as ‘stable’ large gene samples can contain a few significantly upregulated and a few downregulated genes, in effect balancing each other out and not being able to bias a majority stable trend. Thus, we observe 20 genes with significantly upregulated RNA expression under heat shock, of which 12 are annotated as HSPs (Additional file 2: Table S1).

Regarding translation, a similar consistency was also observed amongst the RiboSeq replicas (Fig. 3A-B). Control replicas show Spearman correlation above 0.93 (Fig. 3A), and a near 95% overlap considering ORFs showing RPKM^FP^≥1 (i.e., ≥1 RiboSeq reads, or footprints, per million mapped reads) in either replica (Fig. 3B). We observed a total of 11,692 canonical ORFs with RiboSeq signal above noise (RPKM^FP^ ≥1 in both replicates [24, 42]) (Fig. 3B). Heat shocked RiboSeq replicas show a 0.88 correlation in RPKM^FP^ values (Fig. 3A), and an 86% overlap in canonical ORFs showing RPKM^FP^ ≥1, for a total of 9,953 ORFs with RiboSeq signal (Fig. 3B), 1,739 fewer ORFs than in normal conditions.

**Figure 3.**
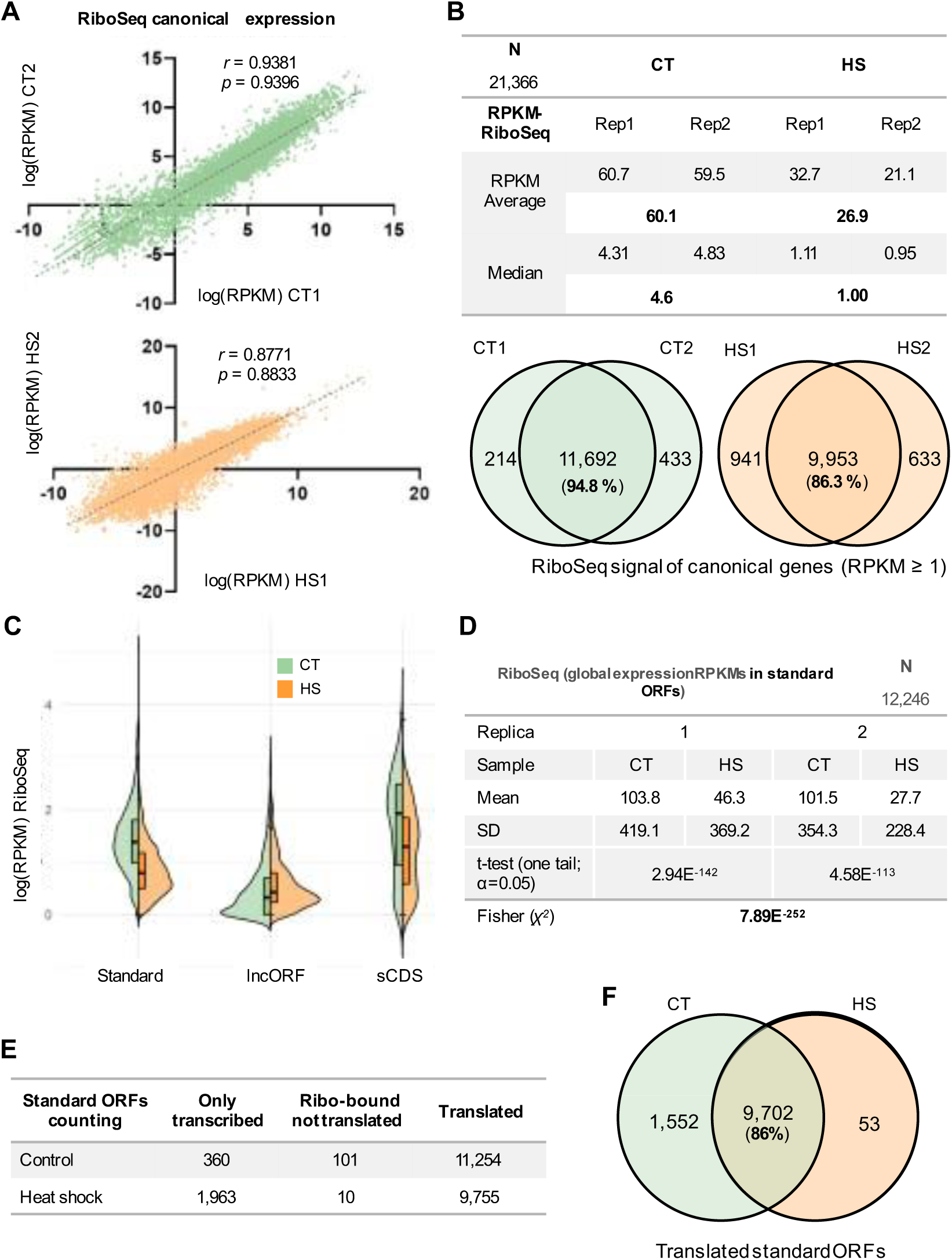
Heat shock reduces canonical ORF translation. **A.** Correlation of RiboSeq signal between replicates for all canonical genes in control (CT, green) and heat shock conditions (HS, orange). (r: Spearman correlation; p: Pearson correlation). **B.** Ribosomal binding signal levels of all canonical genes. The table shows average and median RiboSeq expression per replica and per condition. Venn diagrams show the overlap, between replicas, of genes with RiboSeq signal above noise (RPKMFP ≥ 1). **C.** Distribution of RiboSeq signal across filtered standard ORFs, sCDSs and lncORFs between control and heat shock conditions (see Section “ORF expression analyses in Methods). **D.** Summary of the RiboSeq signal levels of standard ORFs. The table shows mean RiboSeq RPKM values for each condition in each replica, and the results of one-tailed paired t-tests showing significant variation between conditions for both replicas. Overall statistical significance was assessed by Fisher’s method (see Methods). **E.** Summary of the expression status (Only transcribed, Ribo-bound not translated, or Translated) of filtered standard ORFs in control and heat shock conditions. **F.** Overlap of translated standard ORFs between control and heat shock conditions (see Methods and Additional file1: Fig. S2B).

According to the literature, we expected a general reduction in translation as measured by RiboSeq [27, 43] with only genes that provide heat shock resilience (heat shock proteins, or HSPs) showing higher translation [19, 30]. The overall canonical data corroborates this expectation (Fig. 3B, Fig. S2B), but to clarify and quantify these two responses we separated the 21 *Drosophila* ORFs annotated as encoding HSPs from the rest (non-HSP annotated ORFs above 100 aa, henceforth referred to as “standard” ORFs and proteins).

As expected, HSPs show enhanced ribosomal binding (as RPKM^FP^ values) under heat shock, up to several orders of magnitude (Additional file 2: Table S2, Additional file 1: Fig. S3D), confirming that our experimental design induces a bona-fide heat shock. Conversely, standard proteins show an overall and consistent reduction in translation, as shown by a drop in heat shock RPKM^FP^ medians and averages (paired samples one-tailed t-tests with a combined p=7.89E^-252^) (Fig. 3C-D).

By combining RNAseq and RiboSeq the number of actually translated ORFs in either control (CT) or heat shock (HS) can be found, defined as being transcribed, producing ribosomal footprints above background (ribo-bound) and showing codon periodicity (framing) in the ribosomal footprints (p-value < 0.01) (see Methods and [42]). This number is 13.3% lower in heat shock (9,755 vs. 11,254 ORFs) (Fig. 3E-F), and only identifies 53 proteins (0.5% of total translated) as exclusively translated under heat shock (Fig. 3F). A comparison of the translated ORFs reveals that, after 1 h heat shock, 86% (9,702) of standard proteins remain translated (Fig. 3E) albeit at a lower level (as shown above, Fig. 3C-D). This indicates that our heat shock regime has not completely distorted genomic function, avoiding artefactual results.

### smORFs in lncRNAs behave differently to standard genes and maintain or even increase their translation under heat shock

Next, we studied how heat shock affects smORF expression, focusing first on short coding sequences (sCDSs) of less than 100 codons encoded in standard, typically monocistronic, mRNAs (showing polyadenylation and standard-length UTRs). These sCDSs tend to be about 80 codons long, conserved at the class level, and contain one to two structured protein domains (Fig. 1A). These features have facilitated their annotation in *Drosophila*, yet sCDSs are less conserved than the more deeply conserved canonical proteins >100 codons, and encode more hydrophobic and positively-charged amino acids (Fig. 1A, [42]). sCDSs behave similarly to standard ORFs, showing a) robust correlation in replicas in both conditions (Additional file 1: Fig. S3A); b) a general reduction in translation levels under heat shock (Fig. 3C and Additional file 1: Fig. S3D) with only 6% (13 cases) being HS-specific translated (Additional file 1: Fig S3B-C). This is consistent with sCDSs being a class of genes already integrated harmoniously into the functional coding genome, and thus, functionally similar to standard genes, despite coding for shorter proteins and being generally younger (in terms of evolutionary conservation).

ORFs within lncRNAs (lncORFs) behaved differently. First, and as noted previously [24, 42], lncORF translation displays a higher variability than standard genes, showing inter-replica correlations of 0.53 to 0.60 with a 40% to 36% overlap with RiboSeq signal (Fig. 4A-B; Additional file 1: Fig. S3D). Second, contrary to what we observed for standard proteins and sCDSs, we did not observe an average decrease in lncRNA translation, but rather an increase (Fig. 3C, 4C-D). This increase is more acute in one of the replicas, but it is statistically significant overall (paired samples one-tailed t-tests with a combined p=1.24E^-03^, Fig. 4B). A qualitative analysis of translation reveals that this increase extends beyond simple ribosomal binding, to actual productive translation (showing tri-nucleotide framing) (Fig. 4C). Thus, there is an overall increase in the number of translated lncORFs, with 200 lncORFs (56% of total translated lncORFs) newly translated under heat shock (Fig. 4D).

**Figure 4.**
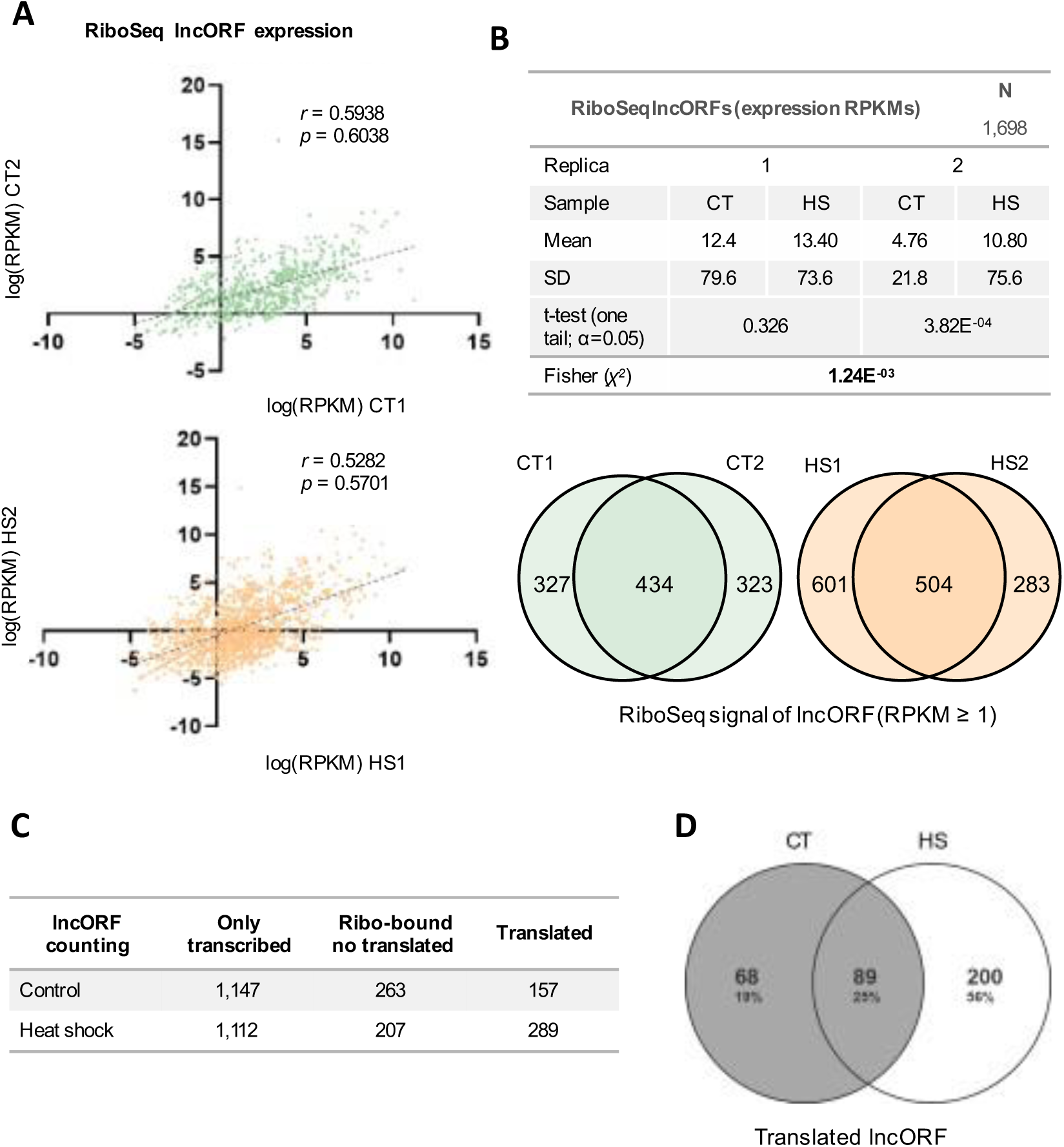
lncORFs remain translated under heat shock. **A.** Correlation of RiboSeq signal between replicates for all lncORFs in control (CT, green) and heat shock conditions (HS, orange) (r: Spearman correlation; p: Pearson correlation). **B.** Summary of lncORFs expression levels. The table shows mean RiboSeq RPKM values for each condition in each replica, and the results of one-tailed paired t-tests on the variation between conditions for both replicas, and their combined significance by Fisher’s method (see Methods). **C.** Summary of the expression status (Only transcribed, Ribo-bound (RPKM ≥1 in both replicas) but not translated (not having significant codon periodicity) or Translated) of filtered lncORFs in control and heat shock conditions. **D.** Overlap of translated lncORFs between control and heat shock conditions according to our pipeline (see Methods Additional file1: Fig. S2B).

### Molecular features of smORF heat shock translation

To explore the molecular features driving this HS-resilient translation, we followed several experimental and analytic avenues. We started by studying monosomes and polysomes separately in one of our samples (RiboSeq replicate 1, Additional file 1: Fig. S2A). Polysomes (formed by several ribosomes bound to a single RNA molecule) arise from intense translation and elongation, whereas monosomes (single ribosomes bound to a single RNA) can result from either new translation initiation, sustained low-level translation, or polysomal degradation. Importantly, translating monosomes (often initiating translation) generate ribosomal footprints (RFPs) with codon periodicity [44] that can thus be distinguished from “silent ribosomes” arising from polysomal degradation [30].

We observe that polysomal RiboSeq signal decreases under heat shock in standard ORFs and sCDSs (Fig. 3C, 5A, Additional file 2: Table S2). The drop in polysomal translation (from near 11,000 translated ORFs with average RPKM^FP^ of 112 to near 8,000 ORFs with RPKM^FP^ of 44) (Fig. 5A and Additional file 2: Table S2) confirms the ‘polysomal collapse’ observed in classical polysomal fractions (Additional file 1: Fig. S1) and hence likely reflects the disassembly of most polysomes following elongation block under heat shock [27]. Moreover, RiboSeq RPKM^FP^ signal in monosomes also decreases for canonicals and sCDS (Additional file 2: Table S2), suggesting a general reduction in all translation-related activities under heat shock.

**Figure 5.**
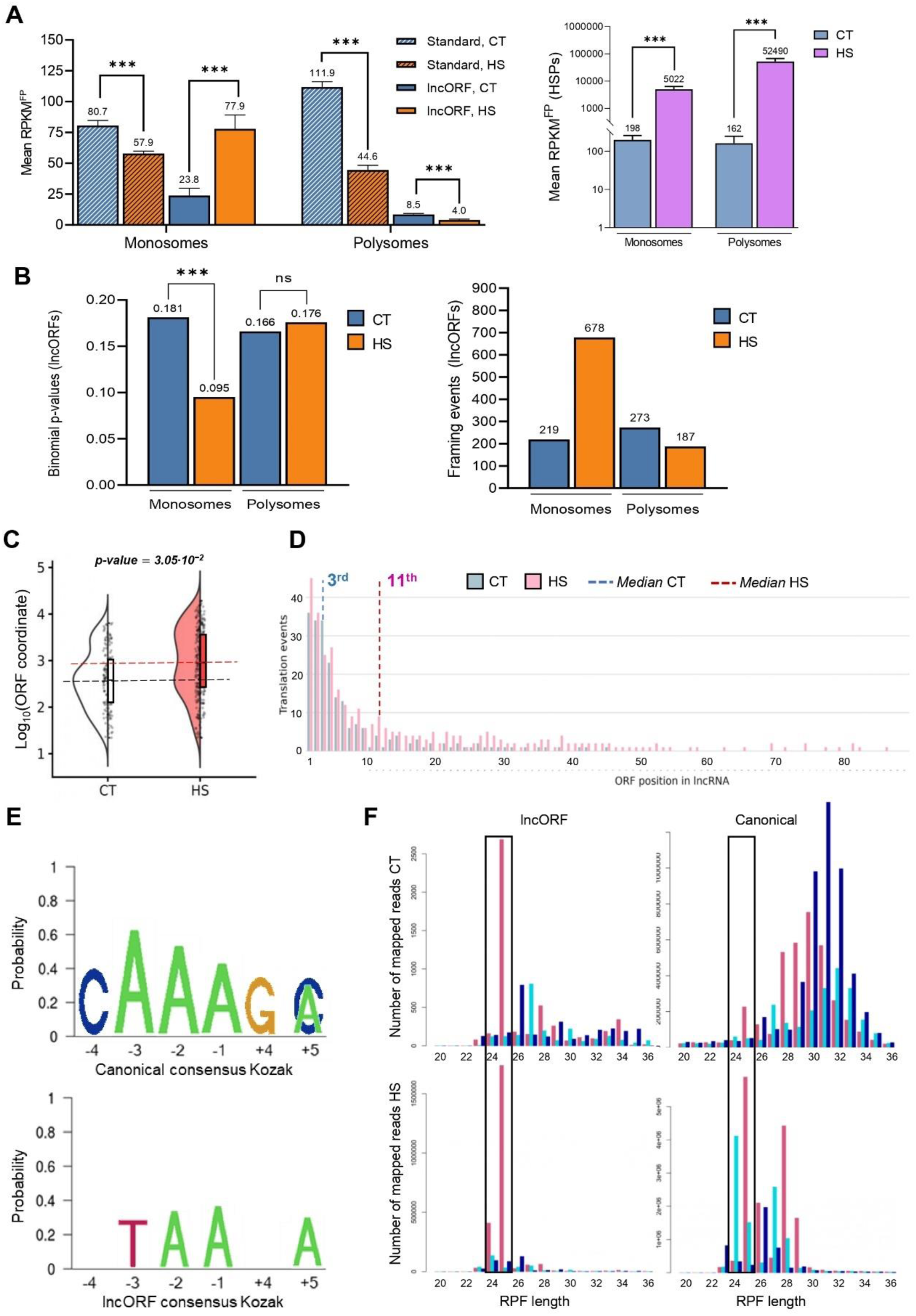
Monosome-mediated lncORF translation, transcript context, and distinctive features of heat-stress translation. **A.** RPKM^FP^ averages found in standard ORFs and lncORFs (left panel) and HSPs (right panel) in monosomal and polysomal fraction in control (blue) and after heat shock (orange/purple). Statistical significance is indicated (t-test p-value *:<0.05; **: <0.01; ***:<0.001; ****: <10^-4^). **B.** Mean of the three-nucleotide framing binomial probability in monosomal and polysomal fractions for translated lncORFs (left panel) and total number of framing events (right panel) (sum of the number of footprint sizes showing framing in each translated lncORF) in lncORFs translated in monosomal and polysomal fractions in control (blue) and heat shock (orange) conditions. Statistical significance is shown as in A). **C.** Absolute position of translated lncORFs in their corresponding transcripts in control (white) and heat shock (red). Y-axis represents the log_10_ of the lncORFs’ nucleotide coordinates from the 5’ end of their lncRNA transcript. Averages are indicated by dotted lines. Statistically significant differences were observed using the Mann-Whitney-Wilcoxon test. **D.** Number of translated ORFs by their relative location to one another lncRNA from the 5’ end (cistronicity). Y-axis: Number of translated ORFs, X-axis: Relative position of the ORF within their lncRNA. Control, CT and heat shock, HS translation are represented in grey and red, respectively. Dashed lines indicate the median cistronic position translated lncORFs in CT (blue) and HS (red). **E.** Kozak consensus sequence for canonical ORFs (top panel) and lncORFs (bottom panel) in *Drosophila melanogaster*, displaying the most representative nucleotides for each position. The Y axis represents the probability of finding each nucleotide at the corresponding position upstream or downstream of position +1, corresponding to the adenine nucleotide of the ATG start codon. **F.** Size distribution of Ribosomal Protected Fragments (RPF; or ribosomal footprint) mapping to translated lncORFs and to canonical genes in control and heat shock conditions. Different colour bars represent the three different frames (red: frame 0; light blue: frame 1; dark blue: frame 2) to which each read maps (see Methods). Box highlights frames in the 24/25 nt footprint sizes.

Contrary to this downward general trend, the results also corroborate the increase of HSPs translation under heat shock. For HSPs, Polysomal RPKM^FP^ increases more than 300-fold, from an already high average expression in controls of about 160 RPKM^FP^, to more than 50,000 under heat shock. Similarly, HSP monosomal RPKM^FP^ increases about 25-fold, from just below 200 to above 5,000 (Fig. 5A right panel, Additional file 2: Table S2). Altogether, the data shows a very strong increase in HSP translation under HS, both at the level of initiation of translation events (monosomes), and at the elongation of these new initiation events plus previously existing ones (polysomes).

lncORFs show more complex results. lncORFs generally display low levels of translation, both in our controls (Fig. 4B-C) and in embryos [42], and this is also shown here with moderate control RPKM^FP^ averages in both monosomes (23.4) and polysomes (8.4) (Fig. 5A left panel, Additional file 2: Table S2). However, the moderate increase in the overall heat shock signal previously noted in replica 1 (Fig. 4B and Additional file 2: Table S2) is observed here to arise from a strong monosomal increase, coupled with a polysomal decrease (Fig. 5A left panel, Additional file 2: Table S2). Thus, monosomal lncORFs signal strength increases significantly about three times, from some 23.8 to 77.9 RPKM^FP^ (Additional file 2: Table S2; paired samples one-tailed t-test p= 3.13^E-06^), whereas polysomal signal is halved from 8.5 to 4.0 (Additional file 2: Table S2; paired samples one-tailed t-test p= 6.11^E-11^). This monosomal increase likely reflects new initiation of lncORF translation. This interpretation is based on an increase in the framing quality of their monosomal signal [44]. Indeed, the average binomial framing p-values for lncORFs in the monosomal fraction lowered from 0.181 in controls to below 0.095 in heat shock (Fig. 5B left panel, Additional file 2: Table S2). Analysing in more detail, we observe that for lncORFs, the number of “framing events” (number of cases where RFPs of different sizes independently show codon periodicity in translated ORFs, see Methods [42]) in the monosomal fraction increases three-fold, from 219 to 678 during heat shock (Fig. 5B right panel, Additional file 2: Table S2), resulting in a net increase in lncORF translation signal by monosomes. Altogether, the analysis of monosomes and polysomes corroborates that lncORFs behave differently to standard proteins, since lncORFs maintain or even increase their translation under heat shock.

We observe a number of molecular features of lncORF translation that diverge from standard translation. One feature is that HS-translated lncORFs have Kozak sequences (which in eukaryotes favour the assembly of ribosomes on translated ORFs [45]) that diverge from canonicity (Fig. 5E). The ‘poor’ Kozak sequences in lncORFs have been noted before and have been linked to the average low levels of lncRNA translation, and their possible inefficient, non-canonical translation mechanisms [11, 42].

Another noticeable feature of lncORF translation is a marked shift in cistronicity (order of the lncORFs in their transcript) to more downstream positions under heat shock. We previously reported that, although translated lncRNAs in embryos are polycistronic and contain several lncORFs, on average those located closer to the lncRNA 5’ are preferentially translated [42]. This 5’ bias for preferential translation of polycistronic lncORFs is reversed during heat shock translation. This is shown, first, by an increase in the average absolute distance of the translated AUGs to the RNA 5’ (Fig. 5C), which is not simply due to a bias for longer lncRNAs being translated under heat shock, since the length of translated lncRNAs is similar in both heat shock (HS) and control (CT) (Additional file 1: Fig. S4A). Second, this is shown by a shift in cistronicity; whereas in controls the median position of translated lncORFs is the 3^rd^ one from 5’ to 3’, and under heat shock is the 11^th^ one (Fig. 5D). This observation is not an artefact of a bias towards polycistronic lncRNAs being preferentially translated under HS, since the number of monocistronic and polycistronic lncRNAs increase similarly (Additional file 1: Fig. S4B). Moreover, the 3’ cistronicity shift is not just qualitative but also quantitative, such as the average RPKM^FP^ signal of lncORFs that were located in the 3’ half (beyond the median position in controls, i.e. 4^th^ or more (Fig. 5D), increases from 16.9 to 45.7 under HS (p= 1.42E-05; Additional file 1: Fig. S4C). This increase in translation levels seems to affect most cistronicity positions along translated lncRNAs (Additional file 1: Fig. S4C). Given that these changes can have functional consequences (see section “Characterisation of *Hsrv*” below), we decided to investigate the 3’ shift in more detail.

We noted that the 3’ shift occurs not only in the 52 lncRNAs that were already translated in controls and continue being translated under HS, which change their cistronicity significantly (“constitutive RNAs”, Additional file 1: Fig. S4D-E), but also when comparing those lncRNAs that are only translated under HS, which have much higher cistronicity than the lncRNAs only translated in controls (“condition-specific RNAs”, Additional file 1: Fig. S4D-E).

One translation initiation mechanism that can produce a 3’ cistronicity shift in “constitutive” lncRNAs is re-initiation, whereby polycistronic mRNAs translate several ORFs sequentially in a 5’ to 3’ direction. This has been shown to occur in several eukaryotic genes [45, 46] including lncRNA-like genes in *Drosophila* [5, 42]. In yeast, stress influences how many and which ORFs are translated in each polycistronic mRNA [45, 47]. The 3’ cistronicity shift in lncORF translation that we observe might be consistent with heat shock favouring re-initiation of translation in polycistronic lncRNAs that were already translating a 5’-located lncORF under normal conditions (Fig. 5C-D). We have used a re-initiation reporter system to study this possibility (Additional file 1: Fig. S4F). *tarsal-less (tal)* is a polycistronic smORF gene expressing several microproteins of 11 to 32 aa, and thus comparable to polycistronic translated lncRNAs [5, 48]. It has been shown that *tal* translation relies on re-initiation [5, 24, 42] under normal conditions, and here, we observe that the translation of a Tal-GFP reporter (Tal2A) is enhanced under heat shock, and that this increase is also mediated by re-initiation (Additional file 1: Fig. S4G).

However, 139 lncORFs are translated only under heat shock from the 51 lncRNAs that showed no translation in controls (Additional file 1: Fig. S4D-E), and therefore these lncORFs cannot benefit from re-initiation. Furthermore, these HS-specific lncORFs show extreme 3’ cistronicity, being located on the 16^th^ position as median (Additional file 1: Fig. S4E), a translation position that has not been reported by any re-initiation studies, and at absolute distances from the 5’ cap (>10^4^, Fig. 5C, Additional file 1: Fig S4E and Additional File 2: Table S4) that might require cap-independent translation [49], suggesting that non-canonical translation mechanisms other than re-initiation may be at work in this context.

Another unusual molecular characteristic of lncORFs is their short ribosomal footprint size. Most ribosomal footprints (RFP) mapping to lncORFs and showing codon periodicity are 24-25 nucleotides (nt) long (Fig. 5F). Interestingly, canonical ORFs (standard ORFs and HSPs) produce two types of footprints: longer, 28-33 nt under normal control conditions, and 24-25 nt ones under heat shock (Fig. 5F). Thus, lncORF footprints resemble canonical ‘stressed’ ones.

Altogether we observe in lncRNAs under heat shock a possible marker of enhanced translational initiation (increased monosomal translation signal) and three molecular characteristics (poor Kozaks, 3’ shift and shorter footprints) that are compatible with non-canonical translation mechanisms (see Discussion).

### *In vivo* relevance of HS-enhanced smORF translation

To clarify the *in vivo* relevance of the HS-enhanced smORF translation, we carried out an RNAseq experiment in adult flies, raised either in normal conditions (controls) or exposed to heat shock. We observe that most S2 cell-translated smORFs are also expressed under the same condition (control or heat shock) in adults (Additional file 1: Fig. S5A-B). Thus, most of the HS-translated smORFs are not S2-specific responses, but more likely universal heat shock responses of the fly cells. This makes sense since, despite instances of insect and dipteran cooling behaviours [50], the heat shock pathway activation is a primary and autonomous cellular response [19, 32, 51]. To further prove *in vivo* relevance, we selected two smORFs for functional characterisation, one sCDS and one lncORF. First, we applied the Z-ratio method (which leverages multiple samples while accounting for their variation, see methods) [52, 53] to identify the smORFs with the highest increase in S2 translation under heat shock. This analysis reveals the significant over-expression of 1 sCDS (*CG6770*) and 12 lncORFs (Additional file 1: Fig. S5C). Second, we cross-referenced them against the lists of smORFs showing heat shock-resilient expression in adult flies (Additional file 1: Fig. S5A-B); and third, we investigated their integration in the heat shock response pathway, whose master switch is the HSF transcription factor (see below). Both the lncORF *heat shock omega 4* (*Hsrv-ORF4)* and the sCDS *CG6770* tested positive for these three criteria. Under heat shock, activation of HSF enhances the expression of HSPs and other genes that increase heat shock resilience (Fig. 1B) [19, 32, 54]. Thus, although overall RNA levels of standard genes and smORFs do not change significantly under heat shock in our samples (see first section of Results), HSF-dependent increased transcription under heat shock is an expected feature of genes directly or indirectly regulated by the heat shock stress pathway [32, 55]. We took advantage of the smORF candidates adult RNA expression and studied their RNA levels by RT-qPCR in flies under control conditions, under heat shock, and under heat shock in mutant flies for HSF (*HSF^4^*, [32]) (see Methods and Fig. 6A). Both the lncORF *Hsrv-ORF4* and the sCDS *CG6770* displayed HSF-dependent increased RNA levels under heat shock, but not when HSF is mutated (*HSF^4^* flies), similarly to the *bona fide* HSF-transcriptional target gene *HSP70* (Fig. 6A), suggesting their direct or indirect regulation by HSF and the heat shock stress pathway. The characterisation of further genes involved in heat shock resilience would be of general interest, and hence we turned to genetics for this purpose.

**Figure 6.**
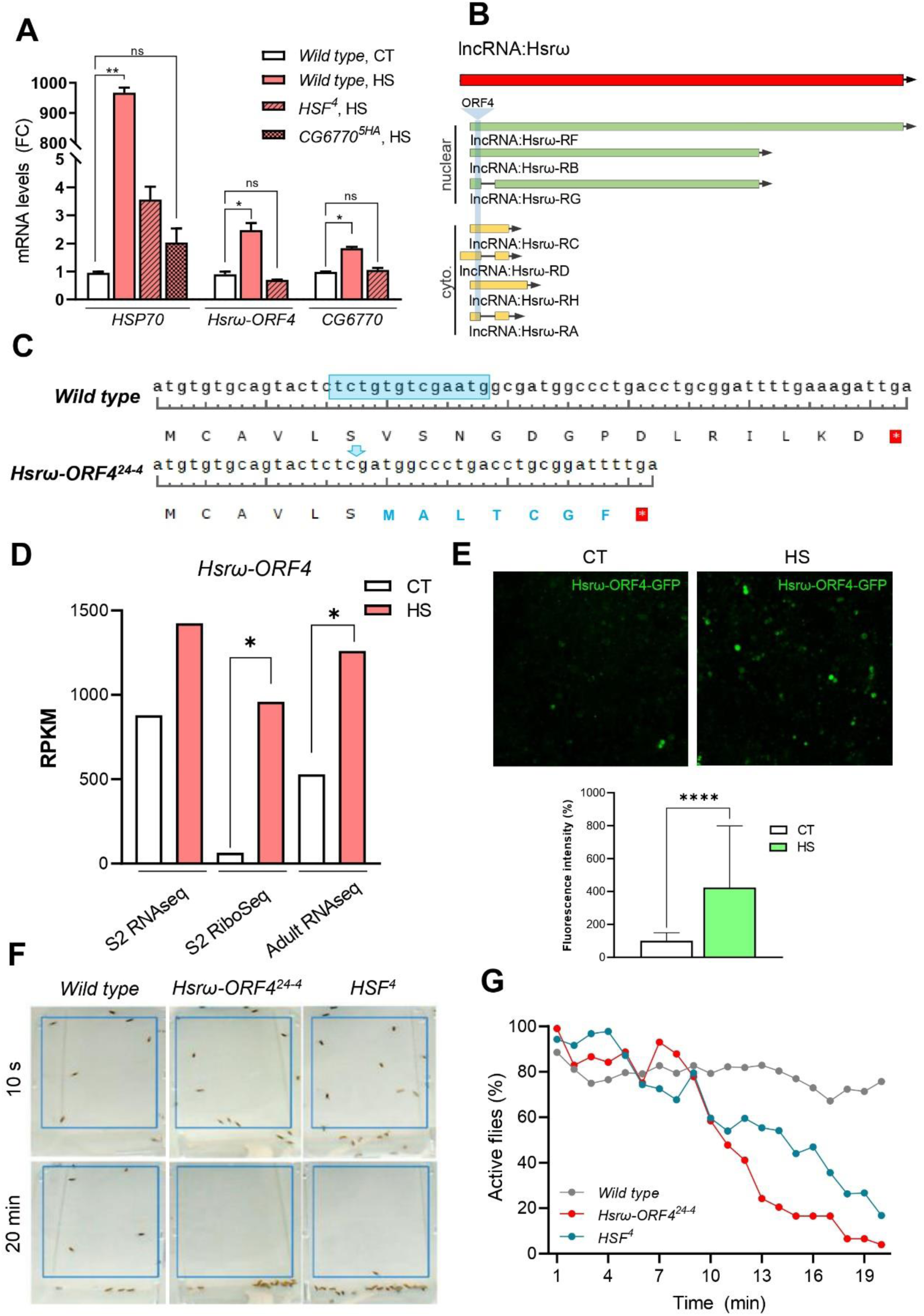
lncORF *Hsω-ORF4* mutant shows strong sensitivity to heat shock. **A.** mRNA levels in control (CT) or heat shock (HS, 1h at 39.5 °C) conditions, as determined by RT-qPCR, for *HSP70*, *CG6770* and *Ηsω-ORF4*, in wild-type, *HSF^4^* and *CG6770^5HA^* adult flies. T-test (two-tailed), ns: non-significant; *p<0.05; **p<0.001. **B.** Schematic representation of known transcripts for *Ηsrω* lncRNA as annotated in Flybase. Green: nuclear transcripts; yellow: cytoplasmic transcripts; light blue: *ORF4* location within *Ηsrω* lncRNA. **C.** Nucleotide and predicted peptide sequences of the wild type and *Ηsω-ORF4^24-4^* alleles generated by CRISPR/Cas9. The *Ηsω-ORF4^24-4^* allele contains a 13-nt deletion that induces a frameshift after the sixth amino acid of the 20aa peptide encoded by HsrωORF4. **D.** RPKM variation of *Ηsrω-ORF4* RNAseq (S2 cells and adult flies) and RiboSeq (S2 cells) samples when comparing CT (control) and HS (heat shock). * Fold change > 2. **E.** Confocal images of S2 cells transfected with Hsrω-ORF4-GFP, in control (CT) or heat shock (HS) conditions (1 h, 37 °C), and quantification of fluorescence intensity. t-test (one tailed), ***p<0.001. **F.** Flies of different genotypes (wild type, *Ηsω-ORF4^24-4^* and *HSF^4^*) were subjected to 39.5 °C heat shock and their activity (number of moving flies) monitored over time. **G.** Percentage of active flies per minute for each genotype. The same triplicated wild type dataset was used as control as in Fig. 6G (see Additional file 1: Fig. S6E).

### Characterisation of *Hsrω*

*Hsrv* is transcribed into several putatively non-coding transcripts (Fig. 6B), whose transcriptional expression has been shown to increase under heat stress [56], which is corroborated by our RNAseq data (Fig. 6D, Additional file 1: Fig. S6A and Additional file 2: Table S1). The longer nucleus-specific RNA isoforms have a non-coding function, associating with and regulating the availability and trafficking of heterogenous nuclear RNA binding proteins (hnRNPs) through RNA-protein complexes known as omega speckles [57, 58]. The shorter isoforms have been detected in cytoplasmic fractions [56], although the function of these cytoplasmic transcripts was not tested. Here we observe that *Hsrv-ORF4* (FBtr0084057_4; 20 codons long) (Fig. 6C and Additional file 1: Fig. S6A), present in the cytoplasmic transcripts of *Hsrv,* is translated (showing high levels of associated ribosomal footprints with significant tri-nucleotide periodicity) and shows a nearly 15-fold increase in translation levels upon heat shock (Fig. 6D, Additional file 1: Fig. S6A and Additional file 2: Table S1). To corroborate the observed increase in translation under heat shock conditions we transfected S2 cells with a C-terminal GFP-tagged version of *Hsrv-ORF4*, obtaining a significant increase in GFP signal under heat shock (Fig. 6E). These results are consistent with the observed increase in RiboSeq footprint signal (Fig. 6D, Additional file 1: Fig. S6A and Additional file 2: Table S1).

We used CRISPR/Cas9 to generate specific mutant alleles in this lncORF with minimal effects on the *Hsrv* transcripts (see Methods). We obtained a mutant *(Hsrv-ORF4^24-4^)* consisting of a 13 nt deletion, that generates a frameshift after the 6^th^ aa of the 20 aa-long peptide encoded by *Hsrv-ORF4* (Fig. 6C). Although this mutation disrupts the encoded peptide, mutant flies were homozygous viable under normal culture conditions (25 °C). Despite this, *Hsrv-ORF4^24-4^* mutants showed a significant reduction in heat shock resilience compared to wild-type. Nearly ten minutes after heat shock, most *Hsrv-ORF4^24-4^* and *HSF^4^*flies succumb to heat shock paralysis, which becomes lethal if prolonged beyond 20 minutes (Fig. 6F-G). This similar behaviour in heat shock resilience and the fact that regulation of *Hsrv-ORF4* is mediated by HSF (Fig. 6A, and section “In vivo relevance of HS-enhanced smORF translation” above), prompted us to corroborate the phenotype of double mutants. *Hsrv-ORF4^24-4^; HSF^4^* double mutants presented a 5 times reduction in viability compared to controls, suggesting that *Hsrv-ORF4* has a role in heat stress resilience downstream of HSF. (Additional file 1: Fig. S6F and Additional file 2: Table S4).

Interestingly, *Hsrv* is a polycistronic lncRNA. Our RiboSeq data and previous reporter assays [56, 59] also indicate the translation of *Hsrv-ORF1*, a lncORF located in the same transcript but 5’ upstream of *Hsrv-ORF4* (Additional file 1: Fig. S6A). RNA expression of *Hsrv-ORF1* also increases under heat shock (in line with the increase observed for *Hsrv-ORF4,* Additional file 1: Fig. S6A, C), but *Hsrv-ORF1* translation levels are not so dramatically upregulated, according to RPKM^FP^ Z-ratios and our S2 reporter system (Additional file 1: Fig. S6A, C-D, Additional file 2: Table S1). Thus, the *Hsrv* gene seems to undergo a quantitative cistronicity shift during heat shock, as the downstream *Hsrv-ORF4* undergoes a relative increase in translation compared to the upstream *Hsrv-ORF1*. We generated CRISPR/Cas9 mutants for *Hsrv-ORF1* (Additional file 1: Fig. S6B) and observed that they neither decrease fly resilience to heat shock (Additional file 1: Fig. S6E) nor display a synergistic interaction with *HSF^4^* mutants (Fig. S6F). Altogether, these results indicate that, contrary to *Hsrv-ORF4*, *Hsrv-ORF1* does not increase its heat shock translation significantly, and has no role in heat shock resilience.

### Characterisation of *CG6770*

The sCDS *CG6770* encodes a 69 aa peptide (FBpp0079890) with predicted alpha helix domains and possible homology to human NUPR1 (Fig. 7A). Although the function of NUPR1 has not been fully characterised, there is evidence for linking it to cancer and stress [60, 61], and for its protein having DNA-binding activity [62]. Accordingly, both NUPR1 and CG6770 contain a nuclear localisation signal (NLS) included in their p8 superfamily domain (Fig. 7B). *CG6770* is translated in control and heat shock conditions, but it shows a 12.4-fold normalised increase in RPKM^FP^ in heat shock compared to control conditions, as well as a 15- to 8-fold increase in RPKM^RNA^ (according to Z-ratios, [53]) (Fig. 7C, Additional file 1: Fig. S3D, S7A; Additional file 2: Table S1). Moreover, CG6770 shows an increase in heat stress-associated shorter footprints (Additional file: Fig. S7B). We followed a similar strategy to that employed when assessing the function of *Hsrv-ORF4*. To test translational regulation by HSF, we introduced *CG6770* minigenes into S2 cells, using an expression plasmid vector whose transcription is not regulated by heat shock. These minigenes contain the endogenous *CG6770* 5’UTR and ORF with an added tag (either GFP or Flag) to allow the detection of the expressed protein ([41], see Methods). We observed a significant increase in CG6770 protein expression under heat shock (Fig. 7D and Additional file 1: Fig. S3D, S7A-C), but this increase was not observed when translation was blocked with cycloheximide (CHX) (Additional file 1: Fig. S7C). These results are in line with the enhanced translation observed with RiboSeq for *CG6770* and corroborate the translational upregulation of *CG6770* by heat shock.

**Figure 7.**
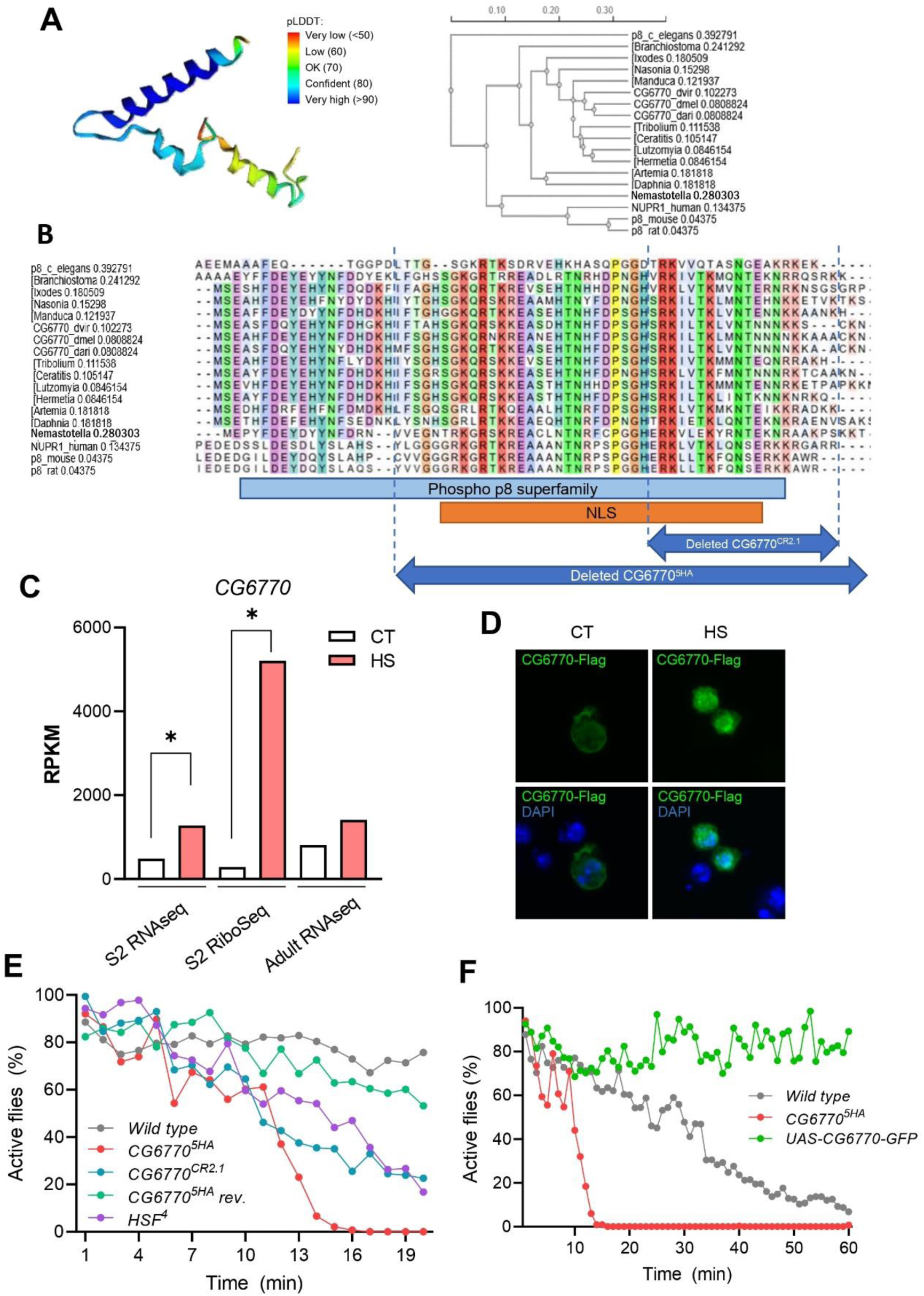
The sCDS CG6770 provides resilence to heat shock. **A.** Left, AlphaFold predicted 3D structure of the CG6770 peptide, coloured by per-residue confidence score (pLDDT). From dark blue regions (pLDDT>90) indicating very high confidence to orange (pLDDT<50) indicating very low confidence (potentially intrinsically disordered regions); right, sequence homology tree of CG6770 and putative homologues, derived by Clustal Omega from the alignment shown in B. **B.** Protein sequence alignment of different *CG6770* homologues employing the “Multiple Sequence Alignment” tool of Clustal Omega software. The consensus Nuclear Localisation Sequence (NLS), Phospho p8 superfamily domain, and the regions deleted in the 5HA and CR2.1 alleles are indicated. **C.** Mean RPKM RNAseq (S2 cells and adult flies) and RiboSeq (S2 cells) values for *CG6770* in control (CT) and heat shock (HS) conditions. * Fold change > 2. **D.** Confocal microscopy images of S2 cells transfected with CG6770-Flag in control (CT) or heat shock (HS) conditions (1 h, 37 °C) (see Additional file 1: Fig S7C). **E, F.** Flies of the indicated genotypes were subjected to 39.5 °C heat shock and their activity (percentage of moving flies) monitored over time. Phenotypes analysed: Wild-type, *CG6770^5HA^*, revertant for 5HA insertion (*CG6770^5HA^ rev.), HSF^4^*, *CG6770* CRISPR/Cas9 deletion mutant (*CG6770^CR2.1^*), CG6770-GFP over-expression mutant (*hs-GAL4/UAS-CG6770-GFP*). In E, the same wild type and *HSF^4^* datasets were used as controls as in Fig. 6G (see methods).

To check the biological relevance of *CG6770* expression and function, we identified a *Drosophila* mutant line (*CG6770^5HA^*) carrying a transposon insertion (P{RS5}CG6770^5-HA-5095^) in the 20^th^ codon of the ORF [63]. This insertion truncates the peptide sequence, deleting most of the Phospho_p8 domain, the NLS and the whole terminal α-helix, and presumably producing a null mutant (Fig. 7B). Even though *CG6770* is highly expressed in control conditions (average RPKM^FP^ = 287), (Fig. 7C) we observed no overt morphological defects nor viability loss in these mutants in normal culture conditions (Additional file 1: Fig. S7D and Additional file 2: Table S3). However, when we submitted these *CG6770^5HA^* mutant flies to heat shock, we observed a dramatic drop in resistance to heat shock compared to control wild-type flies, similar to that observed in flies mutant for *HSF* and *Hsrv-ORF4* (Fig. 7E and Additional file 1: Fig. S7F) [19, 32]. We validated this result in two ways: first, we reverted the mutants by jumping out the (P{RS5}CG6770^5-HA-5095^) insertion (see Methods), which reverted the heat shock resistance to near wild type levels (Fig. 7E); second, we obtained a second mutant allele of *CG6770* by CRISPR (*CG6770^CR2.1^* see Methods and Fig. 7B) and also observed a drop in heat shock resistance (Fig. 7E). Altogether, these results show that the *CG6770* smORF is required *in vivo* for heat shock resilience.

As mentioned above, HSF directs the heat shock response by upregulating the transcription of HSPs (Fig. 1B, 6A). We observed that flies double-heterozygous for *CG6770^5HA^ and Hsf^4^ (CG6770^5HA^/+; Hsf ^4^/+)* are lethal, even at 25 °C (Additional file 1: Fig. S7G and Additional file 2: Table S3). Given that both genes are recessive and viable in heterozygosis (Additional file 2: Table S3; *Hsf ^4^/*+ also reported in Jedlicka *et al*., 1997 [32]), this lethality indicates a strong functional interaction between both genes, compatible with *CG6770* being part of the heat shock response pathway regulated by HSF during heat stress, as with *Hsrv-ORF4*. We next studied *CG6770* effect on the main output of the heat shock response pathway, namely upregulation of *HSP* RNA expression. We observe that, in *CG6770^5HA^* mutants, RNA levels for *HSP70* do not rise significantly under heat shock, indicating that CG6770 is required for *HSP70* RNA upregulation (Fig. 6A).

Finally, to determine whether overexpression of *CG6770* can mitigate the effects of heat shock, we generated transgenic flies carrying the *CG6770* coding sequence fused to GFP, and under the control of the *UAS* promoter (*UAS-CG6670-GFP)* (see Methods). Remarkably, overexpression of *CG6770-GFP* under heat shock conditions (driven by *heat shock* (*hs)*-*Gal4*) (Additional file 1: Fig. S7E), rendered the flies impervious to heat stress (Fig. 7F), showing that *CG6770* is sufficient to provide heat shock resilience, even beyond wild-type levels. Based on its protective function against heat, have named the CG6770 gene as *phoenix (phix)*.

## DISCUSSION

### A robust translational response to heat shock

Our results in cell cultures and *in vivo* recapitulate the heat shock induced translational reprogramming (Fig. 1B and Additional file: Fig S1) [17–20, 32, 51] and add to the corpus of RiboSeq data showing the extent of this reprogramming at the genomic level [14, 27, 43]. In *Drosophila*, 3 h at 37 °C produces a full polysomal collapse, chromosomal puffing and HSP expression, and eventually lethality ([19, 64], unp. obs.). However, flies can recover from a 1h heat shock, indicating that the translational collapse is a regulated, reversible reprogramming. Here we observe that after a 1h heat shock, average translation levels have already declined significantly across the entire standard protein genome, whereas HSP translation is enhanced, altogether supporting heat shock resilience. However, when observed in detail, only a moderate gene fraction has completely lost or significantly reduced their translation, and near 40% of genes remain translated at comparable levels to non-heat-shocked controls. In our system, both up- and down-regulation of individual genes was highly correlated across experimental replicas, suggesting an orderly and predictable regulatory response. This reinforces that translational regulation, albeit not given as much prominence as transcriptional regulation, seems equally robust [24, 25] (this work). We suggest that biological processes that have been characterised as transcriptionally chaotic should be revisited with RiboSeq, in order to ascertain whether stochasticity is a real feature of the process, or merely a transitional feature of transcriptional networks, that is later corrected by translation.

HSPs and other genes that provide resilience to heat stress must avoid the general translational shutdown enforced by heat shock. This can be achieved by translational reprogramming using alternative, non-canonical translation modes that do not rely on the initiation and elongation factors being inactivated by heat stress [26, 65, 66].

Taking framed monosome footprints as indicators of translational initiation events [44], our results suggest that standard protein initiation diminishes during the first hour under heat shock. In parallel, polysome-derived reads reveal a drop in standard protein elongation, observed as lower RPKM, number of ORF translated, number of independent translation events, and loss of polysomal fractions (Fig. 5A, left panel; Fig. 5B, right panel; Additional file 1: Fig. S1). This is compatible with the previously proposed blocks in initiation and elongation under heat shock [26, 27, 29, 43, 67]. Degraded polysomes may convert to monosomes with low codon periodicity (as revealed by the enhanced monosomal fraction, see Fig. 5A-B, Additional file 1: Fig. S1; Additional file 2: Table S2), but freed ribosomes, plus dissociated ribosomal subunits can remain, sometimes associated to stress granules [68]. Thus, a fraction of the free ribosomes and/or ribosomal subunits [21, 30] may bind RNAs, and start the translation of new ORFs by non-canonical mechanisms. HSPs (and some standard genes) could take discretional advantage of this to maintain or even increase their translation levels, perhaps using cap-independent initiation mechanisms such as IRES [69] that bypass the need for eIF2 initiation factor, whose activity is reduced by heat shock and other stresses [26].

### Molecular aspects of lncORF translation under heat shock

smORFs, and especially lncORFs, show resilient and in some cases enhanced translation under heat shock (Fig. 4 and Additional file 1: Fig. S5). A similar observation has recently been reported in maize [36] but to our knowledge this is the first time that this effect is reported in a metazoan. lncORF translation displays features that might be caused by non-canonical mechanisms. Most lncORFs are translated in a polycistronic manner (Fig. S4B), and heat shock induces a 3’ cistronicity shift, such that lncORFs located in more 3’ positions (centered on 11^th^, Fig. 5D) are translated more often and at higher levels. This 3’ bias affects not only lncRNAs that become translated under heat shock but also lncRNAs that were already translated in normal conditions. Thus, not only new lncORFs located in downstream positions become translated under heat shock (revealed by acquiring codon periodicity), but also lncORFs already translated under normal conditions become more actively translated (revealed by their increase in RPKM^FP^) (Additional file 1: Fig. S4C-E). This 3’ shift can result in an increased resilience to heat shock, as in the case of the *Hsrv* gene (see below “Role of smORFs in stress”).

The molecular mechanism mediating this 3’ shift is not fully clear at the moment. It could involve already described mechanisms, new ones, or a combination of both. lncORFs located far from the 5’ end in a polycistronic transcript, must already use non-canonical translation mechanisms (re-initiation-like or IRES-like or other as yet undescribed) when translated under control conditions. These non-canonical mechanisms are not as efficient as cap-mediated ones [70], which may explain the generally low translation levels of smORFs under normal conditions. However, under stress, some of those non-canonical, cap-independent mechanisms seem less affected [20, 22, 23, 28, 29], and this might allow smORF translation to remain at similar levels, and even to increase, taking advantage of freed ribosomes and their subunits [21, 30] similarly as HSPs.

Support for this “freed ribosomes go non-canonical” hypothesis might be provided by the shorter ribosomal footprints produced by lncORFs under both, normal and heat shock conditions (Fig. 5F) [42] but that we also find in HSPs and standard ORFs under heat shock. In the case of HSPs, newly stress-induced translation has been shown to be cap-independent [65, 66, 69, 71] and driven by IRES [69, 71]. It is tempting to speculate that the short footprints (24-25 nt) we detect may be generated by non-canonical translation mechanisms, but given their abundance amongst standard proteins (Fig. 5F), this hypothesis would imply a genome-wide prevalence of non-canonical translation under stress. Alternatively, the difference in footprint sizes (short (24-25 nt) or long (28-30 nt), before and after heat shock, respectively) could be due to different ribosomal elongation states caused by alternative aminoacyl-tRNA incorporation during stress [27, 67]. Under this hypothesis, the prevalence of short footprints in lncORFs could be due to their already non-canonical aa and codon (and hence, tRNA) usage [2, 42], which could delay aminoacyl-tRNA incorporation and elongation [72]. Thus, a heat shock-induced elongation delay may hinder standard ORFs and HSPs, but would not affect lncORFs due to their already low elongation efficiency. Furthermore, smORFs may not need a special elongation mechanism to overcome a stress-induced block. smORF translation could be completed in most cases even if elongation under stress fails around codon 65 [27, 44], since the average translated lncORF size is about 38 codons (this work; [42]). smORF translation might rely on non-canonical mechanisms as a result of chance or necessity [73]. smORFs are mostly young genes [2, 9, 11, 42] and their (generally low) translation under normal conditions might be either accidental noise; or if productive and functional, may not have had the time to evolve high-output mechanisms such as initiation by 5’-cap with eIF2 binding, or efficient Kozak sequences; or elongation by efficient aa and codon usage [2, 42, 74]. Not being dependent on such high-output canonical mechanisms may allow for resilient translation under heat shock, and even increased translation by stress-released ribosomes. This may provide an opportunity for evolution to use these heat-expressed peptides to alleviate stress or help the ongoing genomic translational slowdown. Eventually, some smORFs might be selected to favour stress-expression and function, through optimisation of either their non-canonical translation, peptide activity, or transcriptional regulation by HSF.

### Role of smORFs in stress

Small open reading frames (smORFs) are increasingly recognized as a source of functional peptides with regulatory potential, and genetic assays have repeatedly identified smORF function under stress [3, 4, 12, 14, 35] (this work). Leveraging RiboSeq, RNAseq, and genetics, we have identified stress-related roles for smORFs in the fruit fly. We report widespread translation of smORFs under stress, particularly those located in lncRNAs, in the face of widespread translational slowdown. This persistent, and sometimes enhanced, translation resembles that of genes increasing resilience to heat shock and other stresses, such as HSPs, and hence, suggests a role for smORFs in the response to stress. However, as discussed above, lncORF overproduction might not be biologically significant, but instead a mere accident of these smORFs being shorter, located inside polycistronic lncRNAs, and thus remaining available for non-canonical translation, enduring under heat stress disruption, and taking advantage of increased ribosomal availability. In canonical ORFs, sequence conservation patterns and structural domains can indicate biological function, but these markers are usually absent from smORFs and especially, lncORFs [11, 42]. Hence, one must resort to functional characterisation of a few over-expressed smORFs amongst the hundreds of translated lncORFs and sCDS showing enhanced translation under stress. We have used a two-step approach: first, we identified smORFs with the highest increases in translation (by Z-ratio of RPKM^FP^ [53]). Second, we selected amongst these those also showing an increase in RNA levels (RPKM^RNA^), both in S2 cells and in adult flies, as this makes them candidates for transcriptional regulation by the heat shock pathway mediated by HSF. Thus, these doubly-upregulated smORF candidates are a fair sample of the smORF translational regulation, while their function and biological relevance is easier to demonstrate, by virtue of a possible transcriptional regulation by HSF.

To determine the physiological relevance of *CG6770/phoenix* and *Hsrv-ORF4*, we performed genetic analysis using loss-of-function alleles, which impaired fly resilience to heat shock, and did so in a synergistic manner with *HSF* mutations, suggesting a close functional relationship. Reciprocally, gain-of-function of *phoenix* increased resilience beyond wild-type levels. The relationship of these smORFs with HSF appears to be regulatory, as enhanced RNA expression under heat stress of *phoenix* and *Hsrv-ORF4* is dependent on HSF function. This transcriptional regulation could be direct or indirect, but since HSF is a transcription factor that is activated under heat stress to upregulate the transcription of genes (such as HSPs) that increase resilience to heat stress, the simplest hypothesis is that HSF also upregulates the RNA expression of *phoenix* and *Hsrv-ORF4*. Importantly, we observed that upregulation of HSP70 RNA itself depends on *phoenix*. Given that *phoenix* encodes a putative DNA binding p8 domain [62] and a NLS whose deletion eliminates phoenix function, it is possible that Phoenix acts as a helper of HSF, perhaps increasing HSF transcription factor activity directly or indirectly.

Superimposed to this transcriptional regulation, our RiboSeq and cell culture studies show that *phoenix* and *Hsrv-ORF4* also undergo an increase in translation. This is similar to what happens with HSPs. *Hsrv-ORF4* is a typical HS-enhanced lncORF, having a poor Kozak consensus, and being the 4^th^ ORF in a polycistronic transcript that also translates a microprotein from the 5’ located ORF1. However, the ORF1 peptide seems to play no role in HS resilience (although it may have a function in a yet unknown context), and thus the *Hsrv* gene seems to provide an example for the functional significance of a 3’ translation shift during HS.

The molecular details of the Hsrω-ORF4 and Phoenix microprotein functions are not fully understood at the moment and will require further studies. However, the small 20 aa size of Hsrω-ORF4 microprotein seems to preclude a chaperone, HSP-like activity, but favour a regulatory role on larger, canonical proteins, similarly to previously observed activity for the 29 aa Sarcolamban peptides in muscle contraction [75, 76] and the 11 to 31 aa Tarsal-less peptides during insect leg development [5, 48]. The Phoenix microprotein is larger (69 aa) but similarly sized smORF peptides have also been shown to have regulatory roles [2, 6, 77–79]. Phoenix putative vertebrate homologues (i.e: human NUPR1) have been proposed to have transcription factor-like activity, and to regulate several stress-related proteins [60, 61], and here we observe that *phoenix* is required for *HSP70* RNA upregulation. Regardless of the precise molecular activity of these two smORF microproteins, our results reinforce the view that smORFs play an important and, so far, largely unexplored, role in stress responses. Further, our results suggest a molecular mechanism for the preferential recruitment, and eventual selection, of smORFs towards stress-related functions, based on their non-canonical translation characteristics.

## CONCLUSIONS

1) Small ORFs behave differently under stress than standard ORFs: standard proteins reduce their translation under heat shock whereas small ORFs in lncRNAs maintain it, and in some cases, similarly to HSPs, increase it. 2) The mechanisms that small ORFs use for their polycistronic translation involve a 3’ shift and are likely non-canonical and might be more resistant to the general slowdown of standard translation that occurs during heat shock stress. 3) Two different smORF peptides showing high stress-increased translation are important for conferring organismal-scale resilience to heat shock. The small ORF genes encoding these two peptides are regulated by the heat shock response pathway, showing their integration into a functional genome network.

## MATERIALS AND METHODS

### RiboSeq samples

To subject the S2 cells to heat shock stress, we collected 2×10^8^ cells per experiment and transferred them from the culture flask into a Falcon tube. Control and heat shock samples were then incubated in a water bath for 1 hour at 25°C and 37°C, respectively. Upon heat shock, 5M cells per condition, per replica were kept for total RNAseq. Translation was halted with 100 μg/mL cycloheximide (CHX) (Sigma, St Louis, MO). Cells were pelleted and rinsed twice with 1X PBS supplemented with 100 μg/mL CHX. The pellet was resuspended in lysis buffer [50 mM Tris–HCl pH8, 150 mM NaCl, 10 mM MgCl_2_, 1 mM DTT, 1% NP40, 100 µg/mL cycloheximide, 0.03 U/µLTurbo DNase (Life Technologies, Carlsbad, CA), Complete Protease Inhibitor (Roche)] and incubated on wet ice for 30 minutes. Based on Ingolia *et al*., [80], ribosome footprints were obtained by incubating the lysate with RNAseI (Thermo Scientific™, #EN0601) at 4°C overnight, stopped with SUPERase·In RNAse Inhibitor (Thermo Scientific™, #AM2696), and subjecting the digestion to 4 h ultracentrifugation at 108,000 *g* to pellet all the ribosomal content. 28-35 nt ribosome footprints were purified from a denaturing polyacrylamide gel (25% acrylamide, 19:1 acrylamide:bisacrylamide, 8 M urea). Upon this step, we carried out rRNA depletion with ssDNA, generating by PCR biotinylated anti-sense strings complementary to *Drosophila* 2s, 5s, 5.8s, 18s and 28s rRNA fragments [41]. The resulting oligonucleotides were attached to magnetic Dynabeads^TM^ (Invitrogen^TM^, #65001), and the sense strand was washed away. 50 µL of complementary rRNA beads were incubated during two rounds of depletion to clean the footprints from rRNAs attached to the beads. Purified extracts were then subjected to library preparation by the CABIMER (Centro Andaluz de Biología Molecular y Medicina Regenerativa, Sevilla, Spain) genomics facility. Libraries were then prepared to be sequenced in an NGS Illumina NovaSeq^TM^ 6000 with a 50 base pairs (bp) SingleEnd read protocol. In fractionated RiboSeq experiments, 9 plates S2 cell cultures were grown containing each 50 million cells with 6 plates being heat shocked for 1 h and 3 plates were kept at 25 °C as controls. S2 cells were harvested and lysated with a third of cytoplasmic lysate kept for total RNAseq and the rest loaded onto a sucrose gradient to separate ribosome nucleic acid complexes by centrifugation as described in Aspden *et al*. [41]. Monosome or polysome fractions were extracted using Biocomp fractionator followed by a sucrose cushion step. Ribosome footprint isolation and library preparation and amplification steps were performed as described in Mumtaz *et al*. [81].

Upon sequencing, the NGS facility carried out the adapter and four random base pairs trimming of the reads. We then used Cutadapt [82] to filter by PHRED quality threshold ≥ 33. The remaining rRNAs sequenced in the sample were informatically filtered out aligning the fastq file with a FASTA containing rRNA, snoRNA, tRNA, and snRNA from *D. melanogaster* FlyBase r6.51 genome annotation (dmel_r6.51_FB2023_02) using Bowtie (v. 1.3.1). The non-ribosomal reads were aligned using HISAT2 (v. 2.2.1) [83] with default options to the same *D. melanogaster* genome. We obtained more than 187 million reads aligned to the genome (58 million and 137 million under control and heat shock conditions, respectively) (see Additional file 1: Fig. S2A).

### RNAseq samples

S2 cell cultures for both control and heat shock conditions were derived from the same initial culture described above. Selected flies were grown at 25 °C, and 25 adult males from selected genotypes were subjected to 1 h incubation at either 25 °C (control) or 39.5 °C (heat shock). Total RNA was extracted using the NZY Total RNA Isolation Kit (#MB13402) from approximately 5×10⁶ cells per biological replicate in the case of S2 cells, and 25 adults in the case of flies. Sequencing libraries were prepared by the sequencing facility and subjected to paired-end RNA sequencing of 150 bp fragments. Raw reads were quality filtered using Cutadapt (PHRED quality threshold ≥ 30), yielding a total of 150 and 151 million reads aligned to the genome under heat shock and control conditions, respectively.

### New lncORFs annotation

The new lncORFs list was obtained from the FASTA file downloaded from FlyBase release 6.51 (*dmel-all-ncRNA-r6.51.fasta.gz*). First, we used getorf (EMBOSS-Bioinformatics) to scan from 30 up to 450 nucleotides AUG-STOP regions in the 5’-3’ strand within non-coding transcripts (*getorf -find 1 -reverse No -minsize 30 -maxsize 450*). Thus, a raw count of 25.627 lncORFs is obtained. Second, a bed annotation file is produced, and the lncORFs list is reduced to 18.999 ORFs once duplicated and nested ORFs are cleaned away. Finally, those lncORFs’ coordinates that overlap with canonical, short isoforms, sCDS, uORFs, and dORFs (see classification in Baena-Angulo *et al*. [6] are filtered out using bedtools (*bedtools intersect -v -s -a*) to obtain a list of 18,761 lncORFs. This updated annotation constitutes a substantial improvement over the previous release, adding 2,426 additional IncORFs. Removal of NA and outlier RPKM values leaves the total of lncORFs analysed at 18,492 (see below).

### ORF expression analyses

RNAseq expression levels were quantified as fragments per kilobase of transcript per million mapped reads (FPKM), whereas RiboSeq expression levels were quantified as reads per kilobase of transcript per million mapped reads (RPKM), as RNA-seq and RiboSeq libraries were paired-end and single-end, respectively. FPKM and RPKM values were calculated using the Rsamtools [84] and Seqinr [85] packages in R.

To simplify notation, we refer to FPKM and RPKM values from RNAseq and RiboSeq, respectively, as RPKM^RNA^ and RPKM^FP^ throughout. We removed ORFs with either NA or top outlying RPKM^RNA^ and RPKM^FP^ values (from 10^5^ to ∼2·10^6^) from our pools, to obtain a “raw” total of 21,366 Canonical ORFs, 884 sCDS and 18,492 lncORFs (see “new lncORF annotation” above). For differential expression analyses, we removed ORFs with minimal signal in all samples (RPKM^RNA^=0 or RPKM^FP^<1), to obtain a “filtered” total of 12,267 Canonical ORFs, 277 sCDS and 1,698 lncORFs.

Finally, to identify significantly over-expressed ORFs we used the Z-ratio method [52, 53]: RPKM values were first normalized to Z-scores [(X – sample average)/ SD sample]; then the average Z-score between replicas was calculated [(Zscore Rep1 + Z-score Rep2) / 2], and then the Z-ratio between conditions was calculated (Ave. Z-score CT/ Ave Z-score HS). Following Cheadle *et al*. [53], a ±1.5 Z-ratio threshold was used to obtain ORFs with significantly different expression between conditions.

### ORF’s translation status

Following Patraquim et al. [42], the filtered ORFs were classified as “transcribed” if they obtained RPKM^RNA^ ≥ 1 in any of the two replicas for either CT or HS samples; Ribo-bound, if transcribed and also obtained RPKM^FP^ ≥ 1 in both RiboSeq replicas in either CT or HS samples; and finally, “translated” if Ribo-bound and obtaining a framing probability < 0.01 (Additional file 1: Fig. S2B). To obtain this probability, ribo-bound ORFs must pass a binomial test of framing [42]. We first count the number of total and in-frame reads (see below) in an ORF, separately for each ribosomal footprint size (from 23 to 36 nt) and for each replica and calculate the proportion in-frame reads/total reads. Then we obtain the probability (p) of such proportions in a cumulative binomial distribution with a 1/3 (0.33) random expectation (i.e. randomly, only one in every three FP reads will map in-frame within a given ORF). We take the minimal binomial probability amongst all FP sizes for the studied ORF in each replica, and then combine such minimal binomial probabilities from both replicas using Fisher’s method [86]: a combined test statistic is calculated as ***X*^2^** = −**2** ∑ **ln** (***p_i_***) and evaluated against a ***X*^2^** distribution with four degrees of freedom. We take p<0.01 as statistically significant evidence of framing.

To identify in-frame FP reads, we aligned the non-ribosomal reads (see “RiboSeq samples” above) against the complete ORFome of *D. melanogaster*. This ORFome consists of every AUG-STOP coding sequence in every ORFclass, covering positions from the -18 to +15 nucleotides surrounding the ORF. Using the riboSeqR (v. 1.2.0) R package, each ORF class is analysed separately to assess its global framing pattern. Thus, 23-35 nt RPFs are visualized by bar plots that show the number of reads aligned in frames 0, 1, and 2 at each RPF to assess the three-nucleotide periodicity. Following the same criteria as in Patraquim et al. [42] and the convention in the field, we establish the more abundant frame as the translated one across each ORF class, and then the numbers of total and in-frame reads for each FP size and every ORF are obtained.

### Cell culture and constructs transfection

*Drosophila* S2 cells (S2-DRSC, Stock 181) were sourced from the *Drosophila* Genomics Resource Centre (Indiana, USA). Cells were grown at 25 °C under standard conditions (humidified atmosphere, 5% CO2) in Schneider’s medium with 10% (v/v) FBS and 1% (v/v) penicillin/streptavidin. Transfection was carried out using either pUASt-Hsrω-ORF4-GFP, pUASt-Hsrω-ORF1-GFP, pAWF-CG6770-Flag, pUASt-CG6770-GFP and Act5-Gal4 vectors. The constructs included the 5′ untranslated region (5′ UTR) and coding sequence of the gene of interest, with GFP or the Flag tag fused in frame to the C-terminus of the encoded protein, as appropriate.

### Immunofluorescence

Cells were seeded on glass coverslips, and 48 hours after transfection fixed with 4% pre-warmed paraformaldehyde/PBS for 10 min, permeabilized with 0.1% Triton X-100/PBS for 15 min, blocked with 2% BSA/PBS and incubated overnight with primary antibodies. After being washed with PBS and incubated for 45 minutes with the correspondent secondary antibody and 30 minutes with DAPI, cells were mounted on glass slides with mounting medium (Vectashield Burlingame, CA, USA). Fluorescence images were captured using an SPE Leica confocal laser scanning microscope using a 40×/1.30 oil objective (EC Plan-APO) and LAS AF software. Fluorescence intensity was assessed using ImageJ software.

### *Drosophila* stocks and genetics

Fly stocks and crosses were maintained at 25 °C, unless otherwise stated. Oregon-R (used as *wild-type*) and *HSF^4^* (genotype: *cn^1^ Hsf^4^ bw^1^; P{w[+mC]=Hsf^+t8^}1 / TM3, Sb^1^ Ser^1^*; cat. #5490) lines were obtained from the Drosophila Bloomington Stock Center (BDSC, Indiana University), whereas *CG6770^5HA^*(genotype: *w^1118^; P{RS5}CG6770^5-HA-5095^*; cat. #125824) was obtained from the Kyoto Stock Center. CRISPR guides for the generation of *Hsrv-ORF1^6-12^* (ORF1: GCAGTAGCTACAACCAAAAATGG) and *Hsrv-ORF4^24-4^* (ORF4: GCAGTACTCTCTGTGTCGAATGG) flies were designed using Brown University’s Optimal Target Finder Server (http://targetfinder.flycrispr.neuro.brown.edu/), using the maximum-stringency settings, and cloned into the pCFD1 vector according to the protocol available at http://www.crisprflydesign.org/. The resulting vector was sent for embryo injection to the Centro de Biología Molecular Transgenics Facility, Madrid, Spain, using the *y^1^ v^1^ P{nanos-phiC31\int.NLS}X; P{CaryP}Msp300^attP40^* Drosophila line. *v^+^* transformant F1 females carrying the pCFD1 construct were selected and crossed to *y^1^M{w^+mC^=nos-Cas9.P}ZH-2A w* males. F2 virgin females were collected in bulk and crossed to *w;TM3/TM6B* males. Red-eyed, v+ males were selected and individually crossed to *w;TM3/TM6B* females to establish independent lines (10 lines per ORF). Homozygous flies were used for sequencing of the target locus.

CRISPR/Cas9-mediated genome editing was used to generate the independent *CG6770^CR2.1^* mutant line. Flies carrying a guide RNA targeting *CG6770* from a previously published *Drosophila* line [3] (genotype *y^1^ sc* v^1^ sev^21^; P{y^+t7.7^ v^+t1.8^=TKO.GS01049}attP40/CyO* ; cat. #77235, BDSC, Indiana University) were crossed to flies carrying Cas9 (*y w nos-Cas9*). Progeny carrying the corresponding CRISPR components were selected and used to establish independent lines. Homozygous flies were subsequently used for sequencing of the target locus.

To generate the *CG6770^5HA^* revertant line (*CG6770^5HA^ rev.*), *CG6770^5HA^* flies were crossed to a stock carrying the Δ2-3 transposase (*w; IF/CyO; Δ2-3/TM6B*) to induce excision of the 5HA insertion. Progeny were screened for the appropriate phenotypic and/or genetic markers associated with loss of the insertion, and revertant candidates were individually established as independent lines. The *CG6770* locus was subsequently PCR-amplified and sequenced to identify flies carrying a precise excision event that restores the wild type gene sequence. Candidate lines were further crossed and screened until the *CG6770^5HA^ rev.* allele described in this study was obtained.

To generate *UAS-CG6770-GFP* flies, the pUASt-UAS-CG6770-GFP plasmid was injected into embryos of the 68E attP recipient line (*y^1^ M{vas-int.Dm}ZH-2A w*; M{3xP3-RFP.attP’}ZH-68E*; #24485, BDSC, Indiana University) at the Drosophila Transgenesis Facility of the Centro de Biología Molecular Severo Ochoa (CBM), Madrid, Spain. This line carries the ZH-68E attP landing site on chromosome 3L, allowing targeted integration of the transgene by PhiC31-mediated recombination. Transgenic flies were selected, and independent *UAS-CG6770-GFP*(wt^+^) lines were established. The resulting UAS-CG6770-GFP-containing lines were subsequently crossed to the *hs-Gal4* stock (genotype: *w*; P{w^+mC^=GAL4-Hsp70.PB}89-2-1*; cat. #1799, BDSC, Indiana University) to obtain flies suitable for heat-shock-induced expression of CG6770-GFP.

### Fly motility assays under HS

Before testing, flies were kept at 25 °C. 25 male individuals per genotype, per replica were collected between 7-15 days after pupal eclosion, separated into fly food tubes, grouped by genotype, under CO2 anesthesia, and left to recover for at least 3h at 25 °C. All males from each tube were transferred into T25 cell culture flasks and fully submerged in a water bath at 39.5 °C. Time lapse videos (1 frame every 10 seconds) of the fly-filled flasks were acquired throughout the heat stress assays, using an HD-webcam and the Windows 11 camera app.

Image sequences of each experiment were imported into Image J software, where they were processed as follows: First, a specific ROI was marked for each flask. Second, a lab-developed script was then used in order to count ‘particles’ (flies) per frame, generating a *.tsv archive with the counting data. For each data point, 6 continuous measurements were averaged, to obtain average number of flies in the ROI per minute. All statistical analyses and figures were generated using GraphPad Prism (version 11.0.0 for Windows, GraphPad Software).

### RT-qPCR

Selected flies were grown at 25 °C, and 25 adult males from selected genotypes were subjected to 1 h incubation at either 25 °C (control) or 39.5 °C (heat shock). Total RNA was extracted using the NZY Total RNA Isolation Kit (#MB13402, NZYtech, Lisbon, Portugal). Reverse transcription was performed with the Quantitect Reverse Transcription kit (#205311, Qiagen, Gilden, Germany), and real-time PCR was carried out in a Real-time CFX Connect (Bio-Rad Laboratories, California, USA) with iTaq Universal SYBR Green Supermix (#1725120, Bio-Rad). Amplification efficiencies were validated and normalized against RP49, and fold change in gene expression was calculated using the 2–ΔΔCt method. Results were plotted using GraphPad Prism software. Primer sequences are available in Additional file 1: Table S5.

## DECLARATIONS

### Ethics approval and consent to participate

Not applicable

### Consent for publication

No applicable

### Availability of data and materials

The datasets supporting the conclusions of this article are included within the article (and its additional files).

## Authors’ contributions

CBA, RM, AIP, JIP, ABH and EM performed the experimental work. CBA, RM, AIP, EM and JPC analysed data. RM, JPC and EM secured funding. JPC supervised the work and wrote the first draft. All authors wrote, corrected, read and approved the final manuscript.

## Supporting information

HS movie flies

## Acknowledgements

We thank Sarah Bishop for comments and fly genetics help, Rafael Domínguez Acemel and Alejandro Gil Gálvez for bioinformatics assistance, and Jesús de la Cruz and Juan Tena for discussions.

## Supplementary information

Additional file 1: Figure S1. Polysome fractionation of S2 cells cultured under control and heat shock. Figure S2. Counting of reads in RNAseq and RiboSeq experiments and pipeline outline to determine translated ORFs. Figure S3. sCDSs show canonical-like translation. Figure S4. Features of the 3’ cistronicity shift in lncRNAs. Figure S5. Comparison of smORF RNA expression and translation between control and heat shock in adult flies and S2 cells. Figure S6. The lncORF Hsrω-ORF1 is not involved in the response to heat shock. Figure S7. sCDS CG6770 is essential to survive heat shock. Additional file 2: Table S1 – Expression data.xlsx. Overall RPKM, z-Ratios and translational information. Table S2 – Monosome and polysome data.xlsx. RPKM and p-binomial data in monosomes and polysomes. Table S3 – Drosophila crosses counting.xlsx. Counting of flies in crosses. Table S4 – Cistronicity in lncRNAs.xlsx. Cistronicity analysis, statistical calculations, expression and ORF’s counting in constitutive lncRNAs. Table S5 – qPCR oligos.xlsx. Table with primers used in the qPCR experiments. Additional file 3: HS flies movie.mp4.

## SUPPLEMENTARY FIGURES

**Figure S1.**
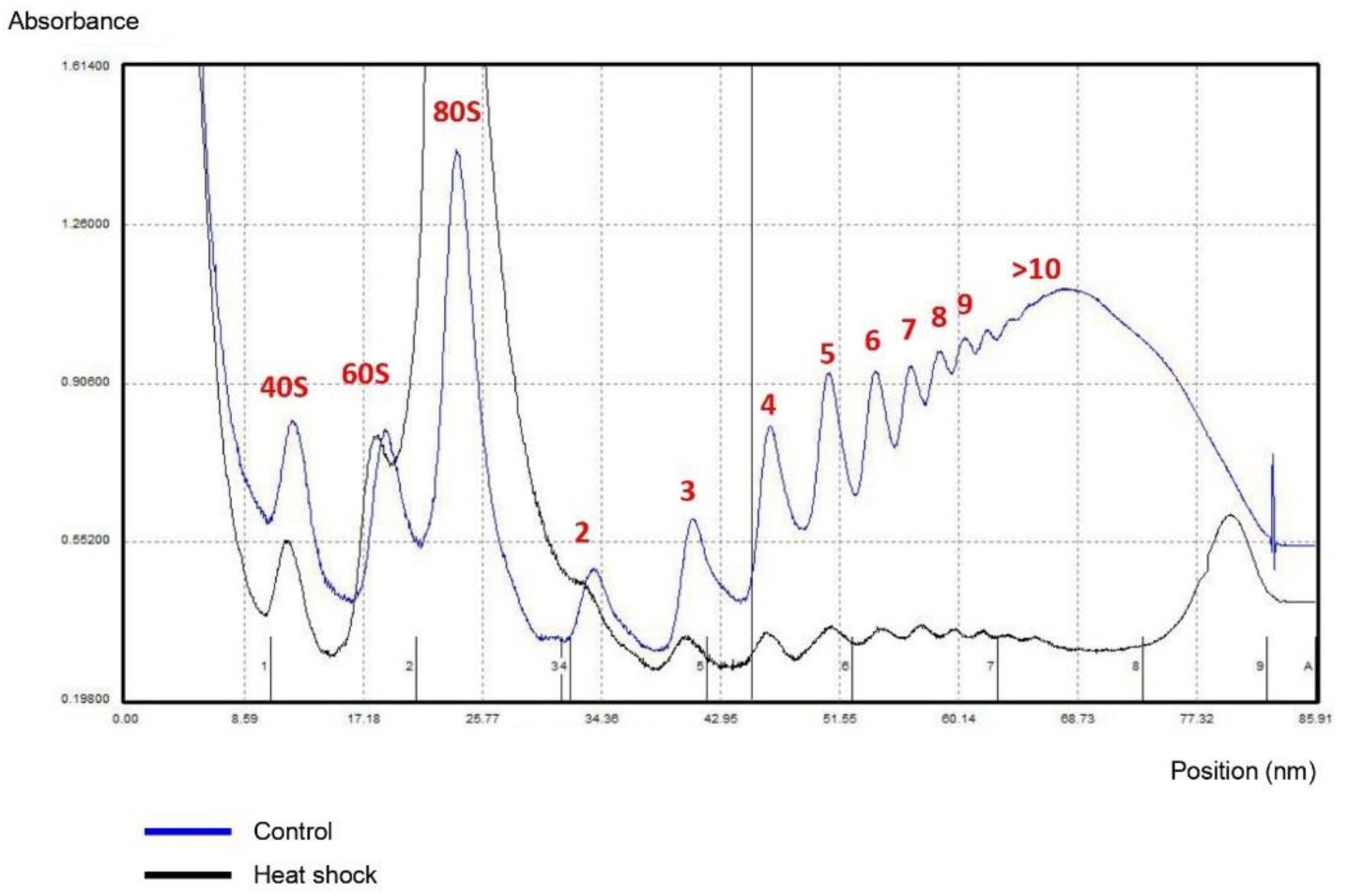
Polysome fractionation of S2 cells cultured under physiological conditions (control) in blue and polysomal collapse after 1 hour of heat shock stimulus in black. Peaks marked as numbers in red from 2 to >10 represents the abundance of polysomal fraction and their number of ribosomes. Peaks marked as 40S, 60S and 80S represent the ribosomal subunit abundance.

**Figure S2.**
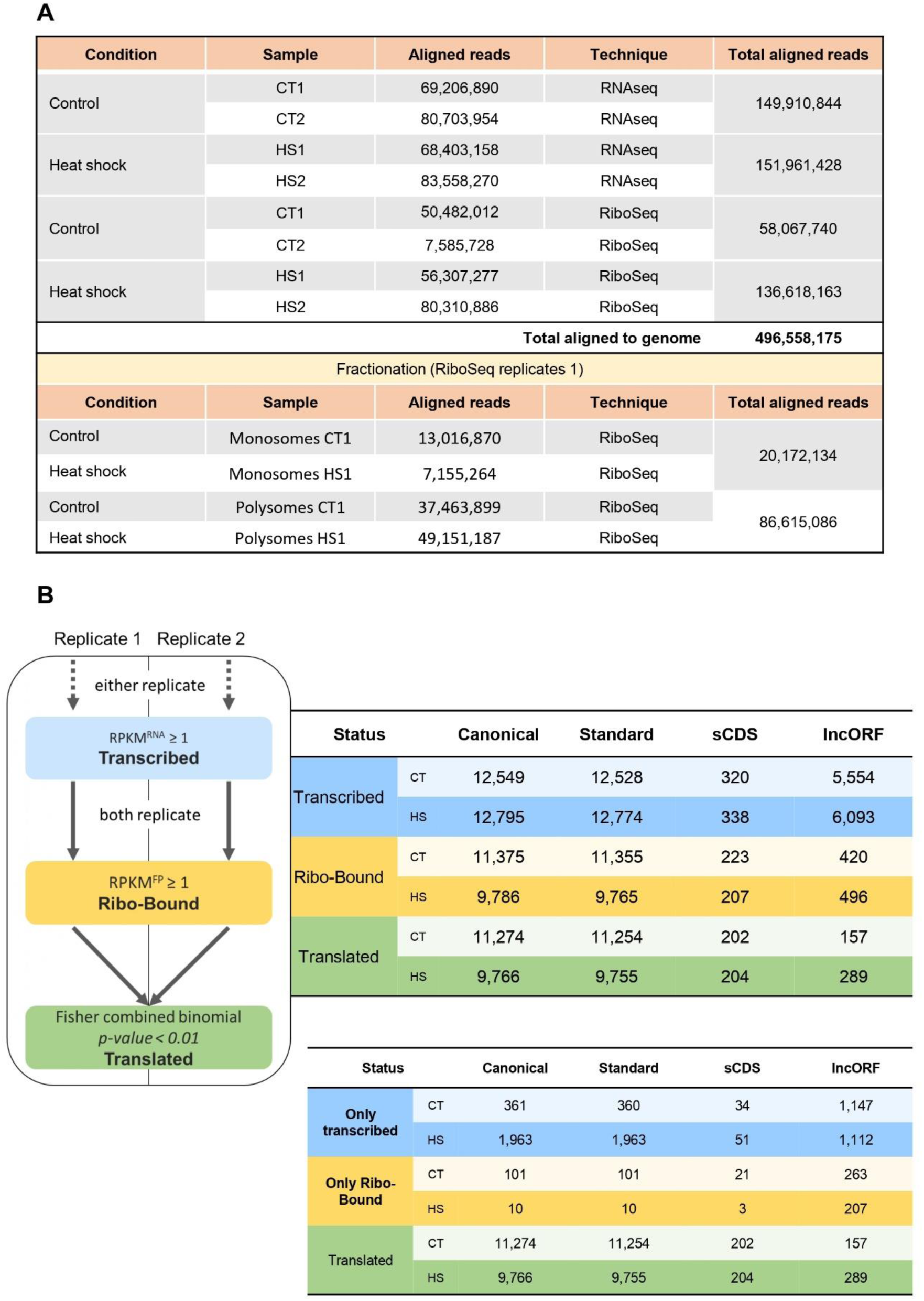
Overall parameters. **A.** Number of non-ribosomal aligned reads to the Dmel Release 6.51 genome in each experiment. **B.** Pipeline to determine the translation status of the ORFs and their respective gene/ORF count. Firstly, to determine the number of translated ORF, we assign the expression status according to three basic levels: Transcribed (blue), Ribo-bound (yellow) and Translated (green). We use the raw population of genes (see Methods: “ORF expression analyses”) and those genes/ORFs with RPKM^RNA^ ≥ 1 in either replicate 1 or 2 was classified as transcribed at their corresponding CT or HS condition. Then, we used the curated list (see Methods: “ORF expression analyses”) to determine “Ribo-bound” or “Translated” states. To be classified as Ribo-bound, those genes classified as transcribed must have RPKM^FP^ ≥ 1 in both replicate 1 and 2. Thus, the transcribed and ribo-bound conditions must also be met in the final cohort of translated ORFs. To finally classify a gene as translated, we look at the binomial probability (*p-binom*) values obtained in the framing analysis of the ribo-bound genes. We used the Fisher’s method (see Methods: ORF’s translation status) to combine the *p-binom* of both replicates and we classify as translated those ORFs with a Fisher *p-value* < 0.01. The table attached to the pipeline outline shows the total number of genes that can be considered as transcribed, ribo-bound and translated in an additive way. Consequently, translated ORFs are also part of the set of ORFs classified as ribo-bound and transcribed. The table below shows the number of genes from the curated gene list that reach only the transcribed, ribo-bound, or translated states.

**Figure S3.**
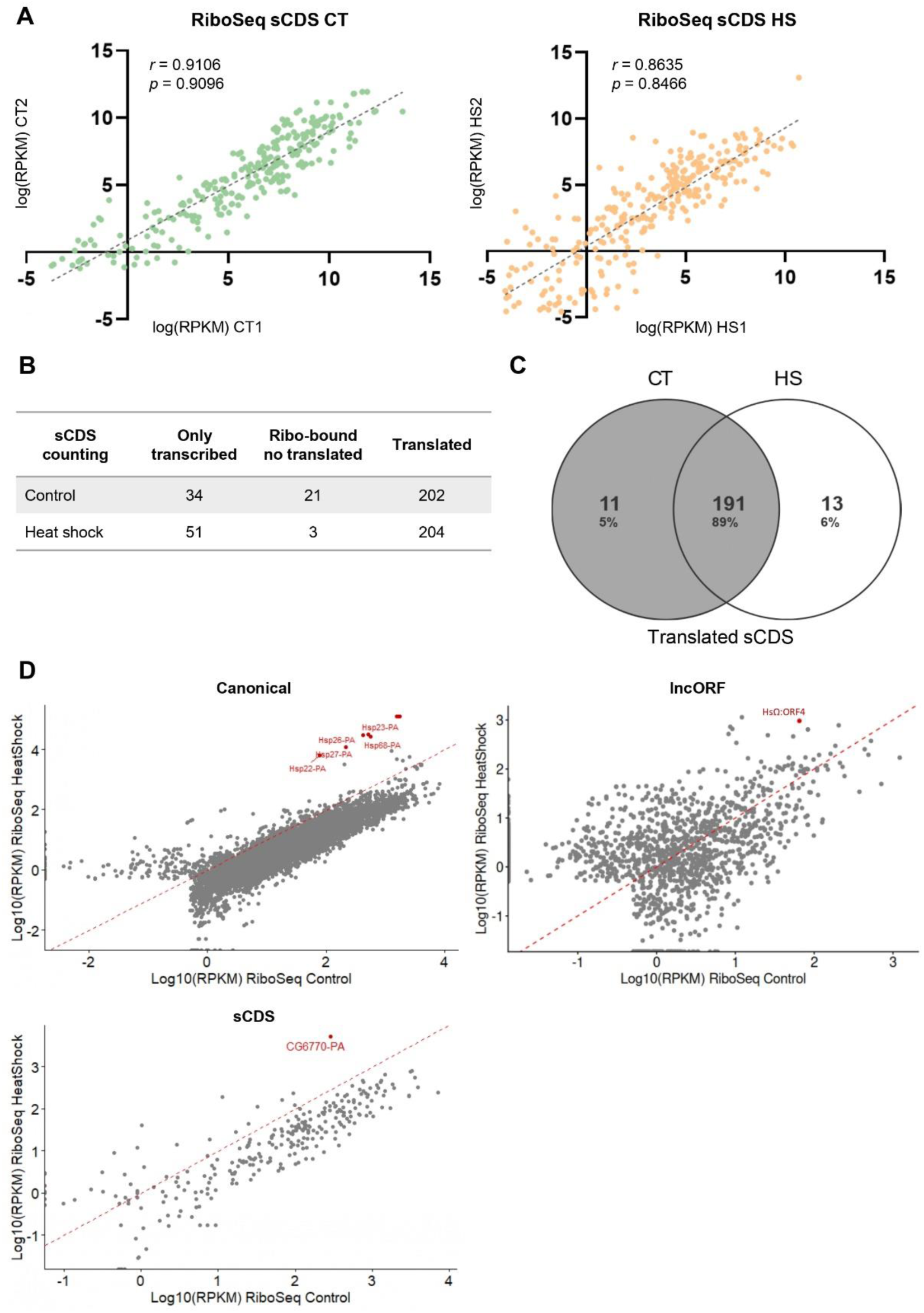
sCDSs show standard protein-like translation. **A.** Correlation of Riboseq signal between replicates for all sCDSs in control (CT, green) and heat shock conditions (HS, orange). (r: Spearman correlation; p: Pearson correlation). **B.** Summary of translational status (Only transcribed, Ribo-bound not translated, or Translated) of sCDSs in control and heat shock according to our pipeline (Additional file 1: Fig S2B). **C.** Overlap of translated sCDSs between control and heat shock conditions. **D.** Dot plot of RiboSeq expression levels between control (X-axis) and heat shock (Y-axis) for canonical ORFs, sCDSs and lncORFs. Plotted RPKMs are the averages between replicates per condition. Up-regulated genes mentioned in text are marked as red dots. Red dashed line follows a slope x=y, indicating unchanged levels.

**Figure S4.**
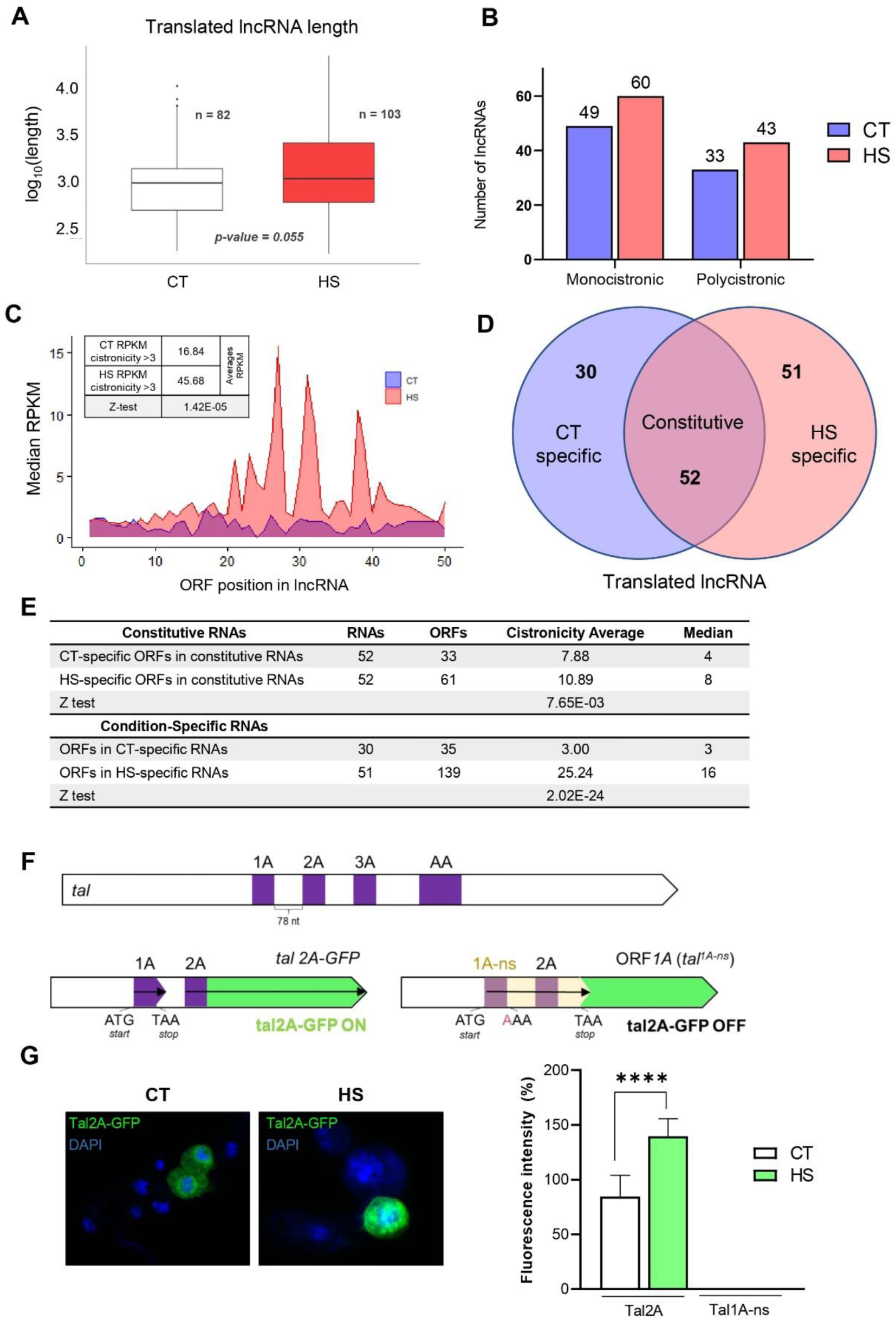
Features of the 3’ cistronicity shift in IncRNAs. **A.** Length comparison of translated lncRNAs in control (white) and heat shock (red) (p-value = U Mann-Whitney-Wilcoxon probability). No significant differences were observed. **B.** Number of monocistronic (only one lncORF translated) and polycistronic (two or more lncORFs translated) lncRNAs translated under control (blue) and heat shock (red). There is a similar linear increase of about +10 lncRNAs in both cases (from 49 to 60 monocistronic to 33 to 43 polycistronic). **C.** Median RPKM^FP^ of the filtered 1,698 lncORFs with RiboSeq signal (see Methods: “ORF expression analyses”) according to their cistronicity position in the lncRNA. Median RPKMs are calculated as the average of the medians obtained in both replicates. Blue, median values for lncORFs expressed under normal conditions; red, lncORFs expressed under heat shock; overlap in purple. Control RPKM^FP^ medians are smaller across most positions on the lncRNAs. The table shows the RPKM^FP^ values of lncORFs located at the third position or beyond under CT and HS conditions. Statistical significance was assessed using 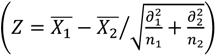 for differences in averages. **D.** Number and type of translated lncRNAs according to their translation in control and heat shock. From left to right: 30 lncRNAs were only translated under control conditions (“CT specific”); 52 were translated under both control and heat shock (“constitutive”) and 51 were only translated under heat shock (“HS specific”). **E.** Average and median cistronicity for control-specific and Heat shock-specific lncRNAs, following panel D: constitutive RNAs significantly increase their cistronicity under heat shock (presumably through changes in their actual lncORFs being translated), and heat shock-specific lncRNAs translate lncORFs with extreme 3’ cistronicity, especially when compared with control-specific lncRNAs. A Z-test of averages was used to compare the means of the two groups and determine whether the observed differences were statistically significant. **F.** The *tal* ORFs are translated by re-initiation from non-overlapping smORFs arranged in different frames from 5’ to 3’ (purple boxes in top *tal* gene diagram). Two different constructs assess the re-initiation of the tal-A2 ORF (bottom diagrams, [42]). The tal2A-GFP construct (bottom left) contains the native 5’ half of the *tal* mRNA, including the tal1A and tal2A microprotein ORFs (purple) tagged with GFP (green) at its C-terminal. The tal1A-ns construct (bottom right) has a mutation in the 1A STOP codon that extends the tal1A ORF (tal1A-ns, arrow) over and beyond the tal2A AUG start. In normal conditions, GFP expression is produced by the tal2A-GFP construct (see panel G below, [5, 42]), and in principle this Tal2-GFP translation could be driven by either re-initiation or an IRES. However, the lack of GFP translation when the tal1A ORF is extended over the tal2A start in the tal1A-ns construct (panel G, below and [42]) indicates that tal2A translation is driven by re-initiation, as reinitiating scanning ribosomes cannot “walk backwards” 5’ towards the tal2A start after terminating tal1A-ns translation 3’ downstream [45, 47]. Conversely, IRES-mediated translation of tal2A could still act in the tal1A-ns construct, but it does not, discarding that IRES-like mechanisms mediate tal2A translation. **G.** S2 cells transfected with Tal2A-GFP or Tal1A-ns under control and heat shock conditions (1 h, 37 °C). GFP expression of the tal2A-GFP construct is observed under control, and it is significantly increased under heat shock (images, left, and quantification, right). However, GFP is not observed in cells carrying the tal1A-ns construct, either under control or heat shock. Altogether these results show that polycistronic translation by re-initiation occurs in S2 cells and that it can be enhanced by heat shock. Cells were analyzed by confocal microscopy, and fluorescence intensity was quantified for each condition. ****p<10^-4^.

**Figure S5.**
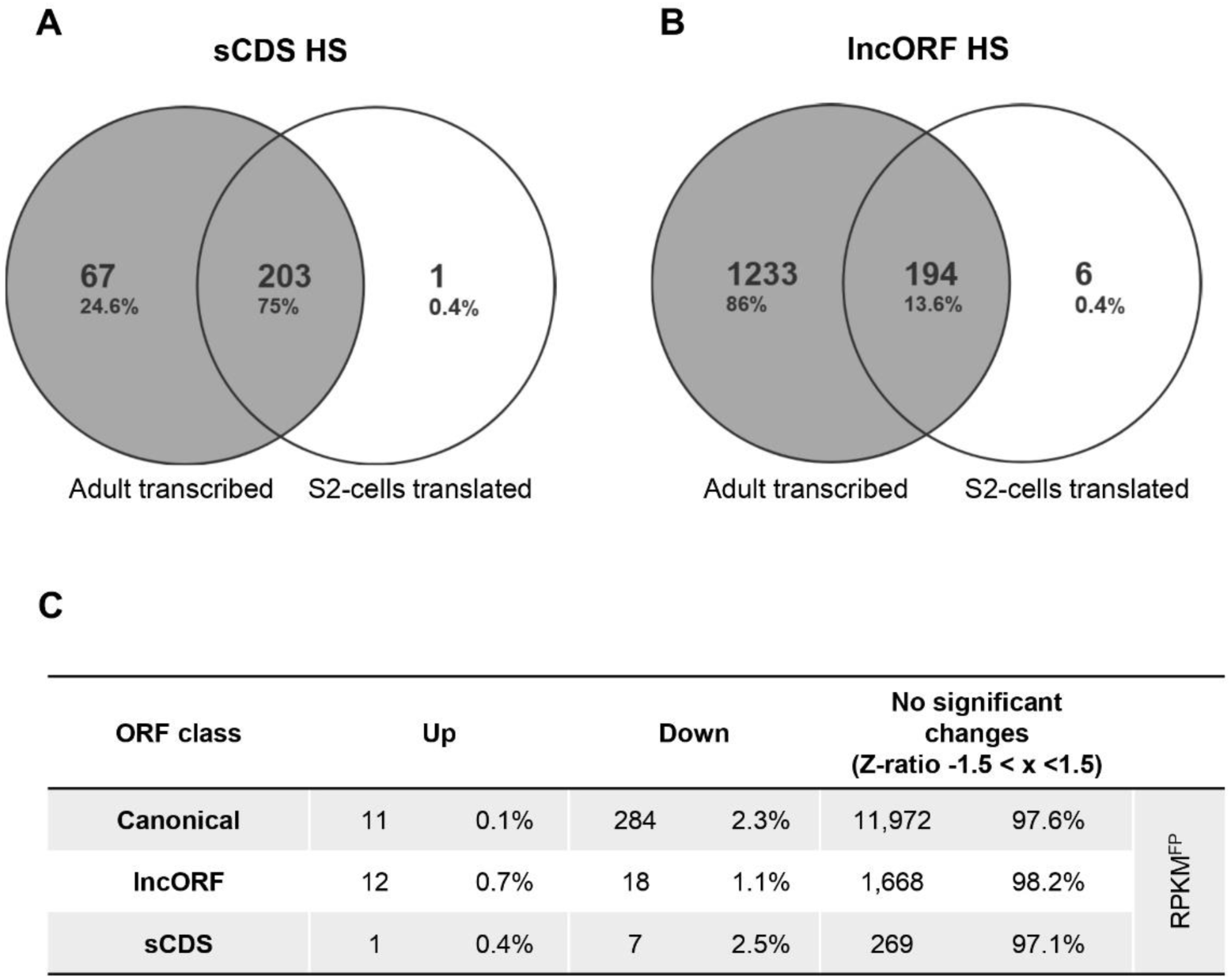
Comparison of smORF RNA expression and translation between control and heat shock in adult flies and S2 cells. **A.** Overlap between sCDS showing RNA expression under heat shock in adult flies (RPKM^RNA^ ≥ 1) and heat shock-translated sCDSs in S2 cells. Almost all S2-translated sCDS are also expressed in adult flies. **B.** Overlap between lncORFs with RNA expression under heat shock in adult flies (RPKM^RNA^ ≥ 1) and heat shock-translated lncORFs in S2 cells. Most S2-translated lncORFs are also expressed in adult flies. **C**. Differentially translated genes across ORF classes in response to heat shock. Number and percentage of canonical ORFs, lncORFs, and sCDSs that were upregulated, downregulated, or showed no normalised significant changes under heat shock (HS) compared with control conditions (CT). RPKM^FP^ translation levels were normalized and their differences assessed by their Z-ratio fold change between control and heat shock (see Methods: “ORF expression analyses”).

**Figure S6.**
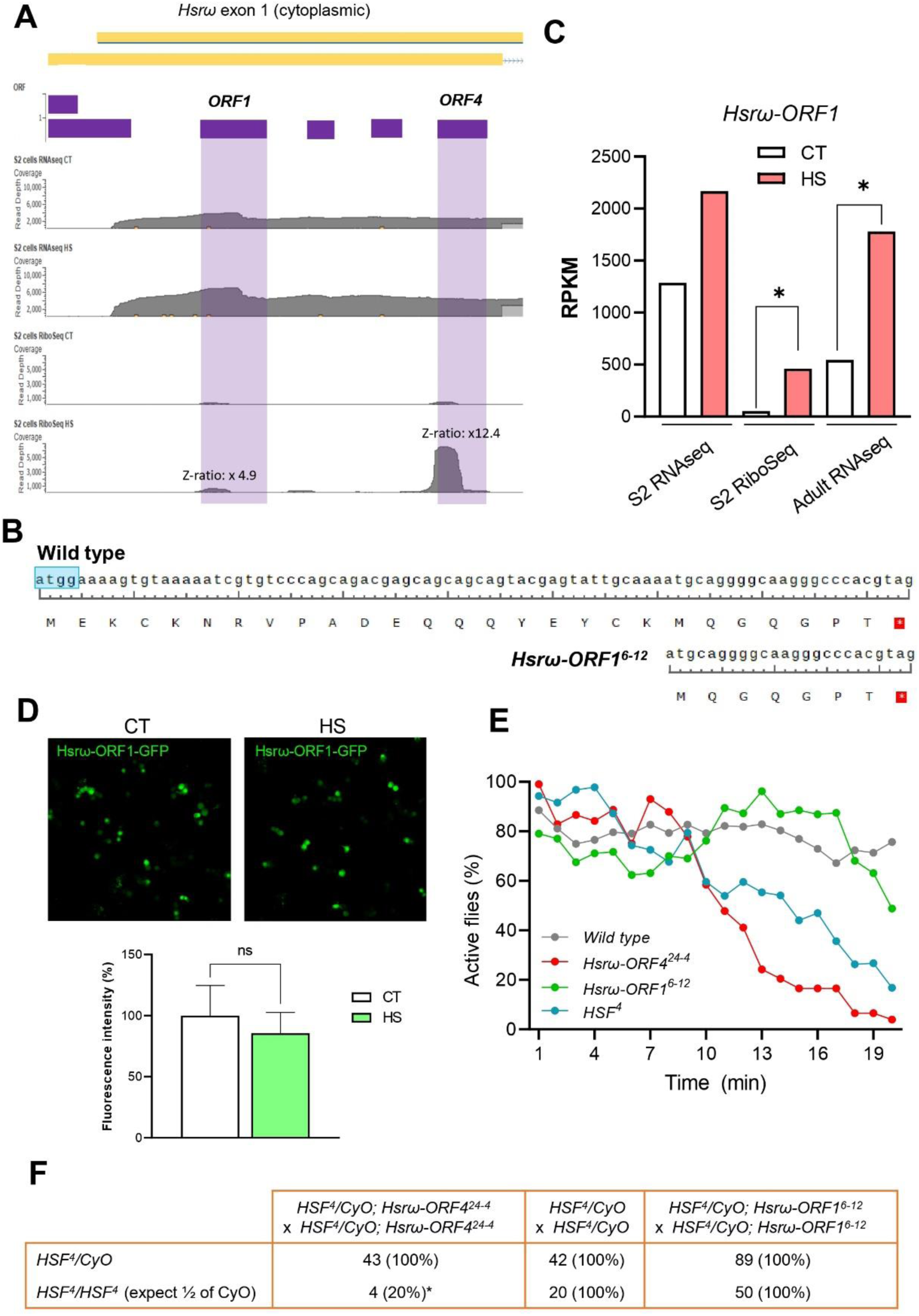
The lncORF *Hsω-ORF1* is not involved in the response to heat shock. **A.** RNAseq (upper panels) and RiboSeq (lower panels) read density map for the *Ηsrω* exon 1 (yellow bars), in both CT and HS. *ORF1* and *ORF4* location is highlighted (shaded in purple). The Z-ratio changes for Ribo-seq are indicated. **B.** Nucleotide and predicted peptide sequences of the *Ηsrω-ORF1^6-12^* allele generated by CRISPR/Cas9. This allele contains a 12-nt deletion that eliminates the start codon of the 27-amino-acid peptide encoded by Hsrω ORF1. If translation were to initiate at the downstream methionine at position 20, the resulting peptide would consist of only 7 amino acids. **C.** RPKM variation of *Hsrω-ORF1* RNAseq (S2 cells and adult flies) and RiboSeq (S2 cells) samples when comparing CT (control) and HS (heat shock). * Fold change > 2. **D.** Confocal microscopy images of S2 cells transfected with Hsrω-ORF1-GFP in control (CT) or heat shock (HS) conditions (1 h, 37 °C), and quantification of fluorescence intensity showing no differences in GFP expression between these conditions. t-test (one tail), ns: non-significant. **E.** Percentage of active flies over time for each indicated genotype, while flies were subjected to heat shock. The same wild-type and *HSF^4^* datasets were used as controls across experiments (see Fig. 6G and methods). **F.** The viability of *HSF^4^* homozygous mutant flies was assessed in the presence or absence of *Hsω-ORF4^24-4^* and *Hsω-ORF1^6-12^* mutant alleles and compared with the viability of heterozygous *HSF^4^* /+ as controls. Raw numbers and percentages from expected ratios are shown.

**Figure S7.**
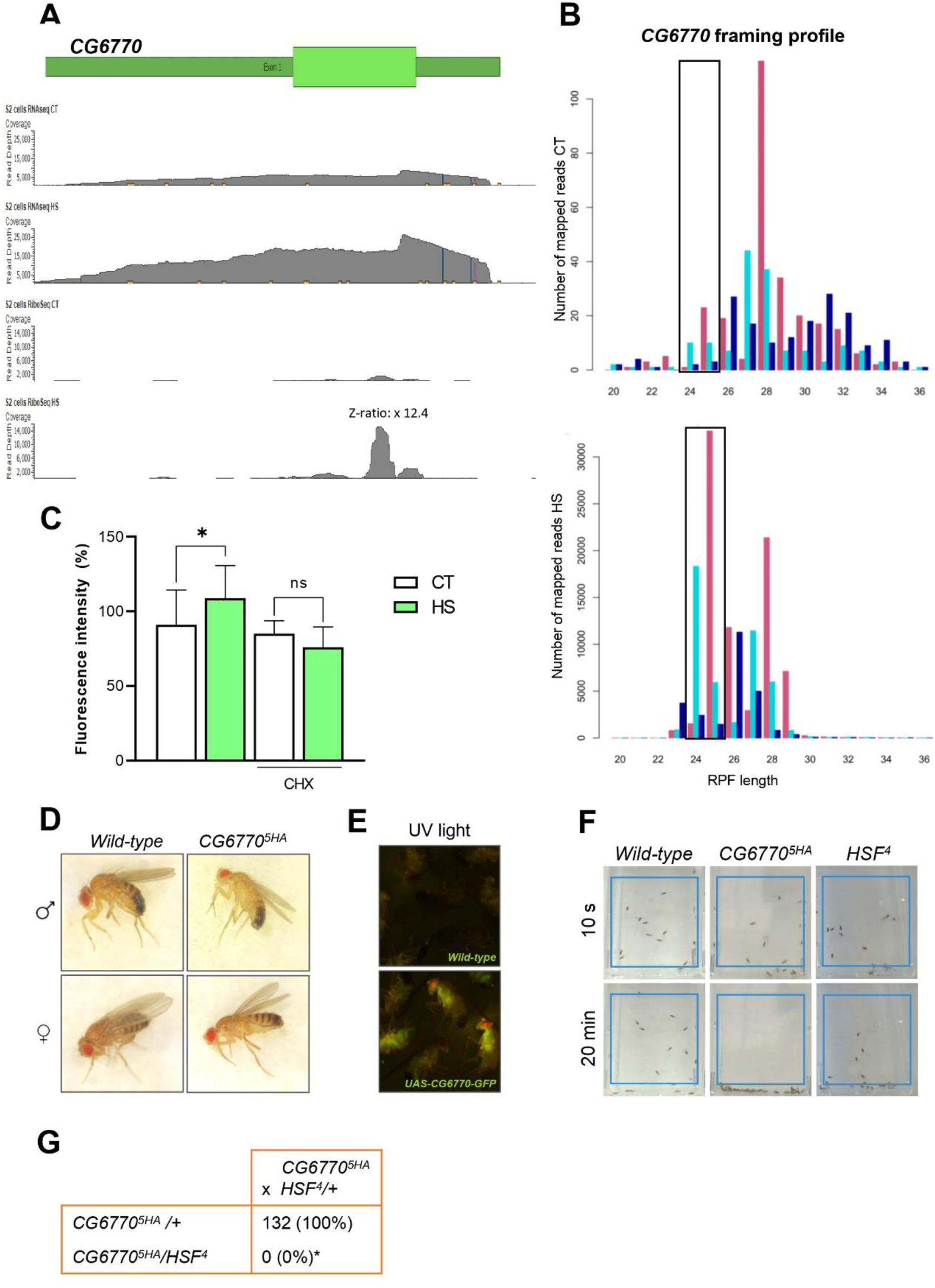
sCDS *CG6770* is essential to survive heat shock. **A.** RNAseq (upper panels) and RiboSeq (lower panels) read density map for the *CG6770* locus, in both CT and HS conditions. **B.** RPF size distribution of *CG6770* in control (CT) and heat shock (HS) conditions. Different colours represent reads aligned to different frames (red: frame 0; green: frame 1, blue: frame 2). Box highlights frames in the 24/25 nt size footprints**. C.** Quantification of fluorescence intensity for CG6770-GFP expression in S2 cells in CT or after 1h heat shock, with or without cyclohexamide (CHX) treatment. ns: non-significant; t-test (one tail), *p-value<0.05. **D.** Images of wild type and *CG6770^5HA^* flies, male and female, showing no gross morphological abnormalities. **E.** CG6770-GFP (*hs-GAL4/UAS-CG6770-GFP*) flies imaged under UV light after heat shock treatment, showing CG6770-GFP expression. **F.** Wild-type, *CG6770^5HA^* and *HSF^4^* flies were submitted to 39.5 °C incubation and their activity monitored over time. Example of some photograms are shown (10 s and 20 min into the experiment). **G.** *CG6770^5HA^* flies were crossed with *HSF^4^* and the F1 progeny was counted. Raw numbers and percentages from expected mendelian ratios are shown.

