## Supplementary figures for "Translation of small ORFs in non-canonical RNAs provides resilience to heat shock stress": Fig_S2.pdf

**A**

| Condition | Sample | Aligned reads | Technique | Total aligned reads |
|---|---|---|---|---|
| Control | CT1 | 69,206,890 | RNAseq | 149,910,844 |
| | CT2 | 80,703,954 | RNAseq | |
| Heat shock | HS1 | 68,403,158 | RNAseq | 151,961,428 |
| | HS2 | 83,558,270 | RNAseq | |
| Control | CT1 | 50,482,012 | RiboSeq | 58,067,740 |
| | CT2 | 7,585,728 | RiboSeq | |
| Heat shock | HS1 | 56,307,277 | RiboSeq | 136,618,163 |
| | HS2 | 80,310,886 | RiboSeq | |
| | | | **Total aligned to genome** | **496,558,175** |
| Fractionation (RiboSeq replicates 1) | | | | |
| **Condition** | **Sample** | **Aligned reads** | **Technique** | **Total aligned reads** |
| Control | Monosomes CT1 | 13,016,870 | RiboSeq | 20,172,134 |
| Heat shock | Monosomes HS1 | 7,155,264 | RiboSeq | |
| Control | Polysomes CT1 | 37,463,899 | RiboSeq | 86,615,086 |
| Heat shock | Polysomes HS1 | 49,151,187 | RiboSeq | |

**B**

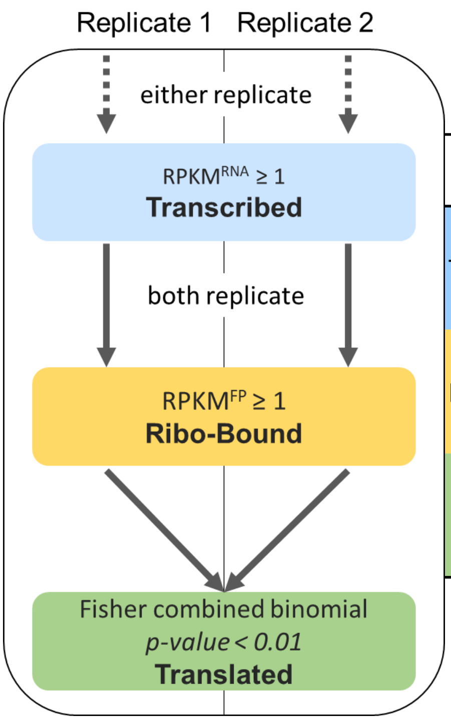


| Status | | Canonical | Standard | sCDS | lncORF |
|---|---|---|---|---|---|
| Transcribed | CT | 12,549 | 12,528 | 320 | 5,554 |
| | HS | 12,795 | 12,774 | 338 | 6,093 |
| Ribo-Bound | CT | 11,375 | 11,355 | 223 | 420 |
| | HS | 9,786 | 9,765 | 207 | 496 |
| Translated | CT | 11,274 | 11,254 | 202 | 157 |
| | HS | 9,766 | 9,755 | 204 | 289 |

| Status | | Canonical | Standard | sCDS | lncORF |
|---|---|---|---|---|---|
| **Only transcribed** | CT | 361 | 360 | 34 | 1,147 |
| | HS | 1,963 | 1,963 | 51 | 1,112 |
| **Only Ribo-Bound** | CT | 101 | 101 | 21 | 263 |
| | HS | 10 | 10 | 3 | 207 |
| Translated | CT | 11,274 | 11,254 | 202 | 157 |
| | HS | 9,766 | 9,755 | 204 | 289 |
