## Supplementary figures for "Translation of small ORFs in non-canonical RNAs provides resilience to heat shock stress": Fig_S4.pdf

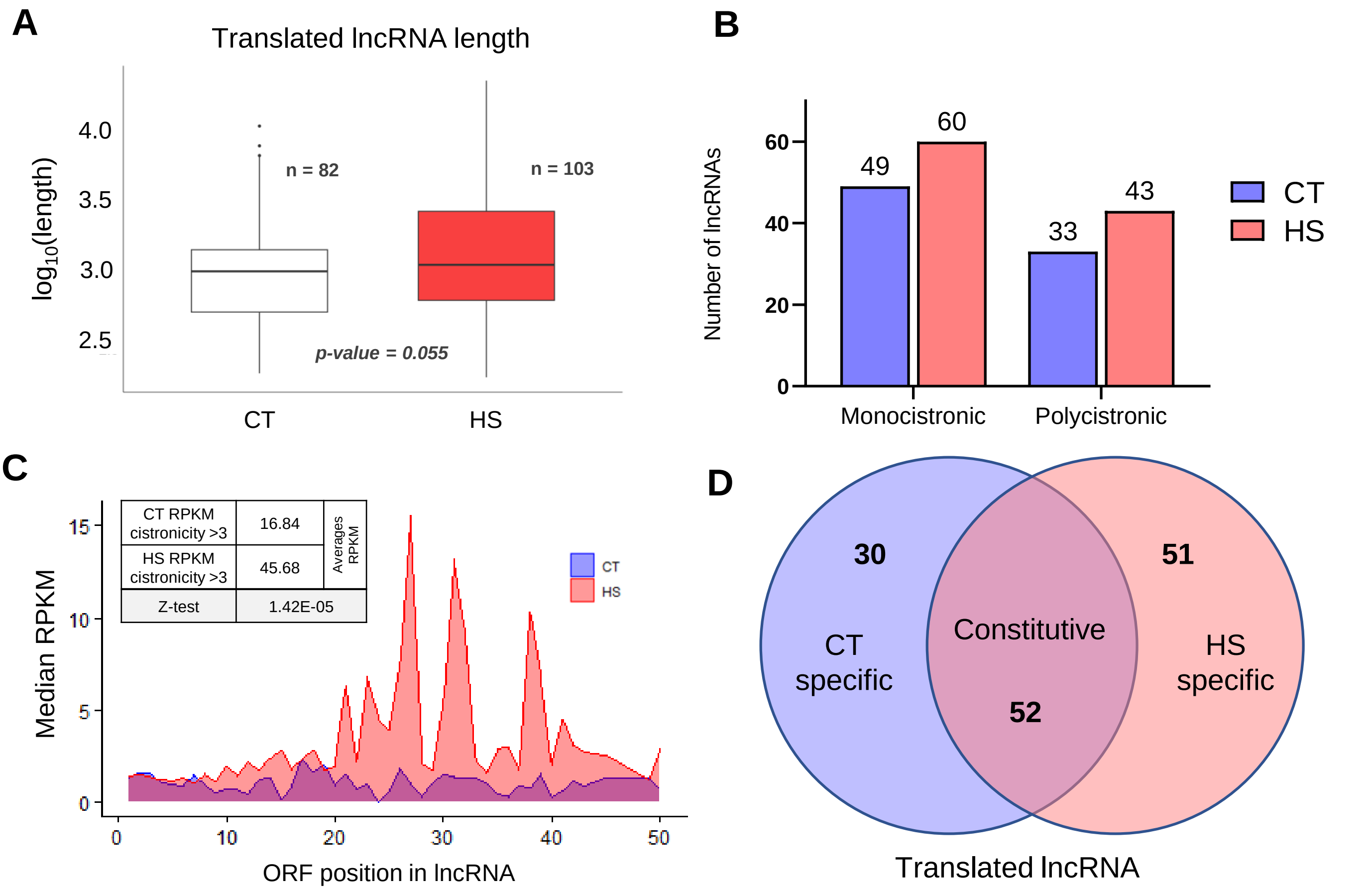


**E**

| Constitutive RNAs | RNAs | ORFs | Cistronicity Average | Median |
|---|---|---|---|---|
| CT-specific ORFs in constitutive RNAs | 52 | 33 | 7.88 | 4 |
| HS-specific ORFs in constitutive RNAs | 52 | 61 | 10.89 | 8 |
| Z test | | | 7.65E-03 | |
| **Condition-Specific RNAs** | | | | |
| ORFs in CT-specific RNAs | 30 | 35 | 3.00 | 3 |
| ORFs in HS-specific RNAs | 51 | 139 | 25.24 | 16 |
| Z test | | | 2.02E-24 | |

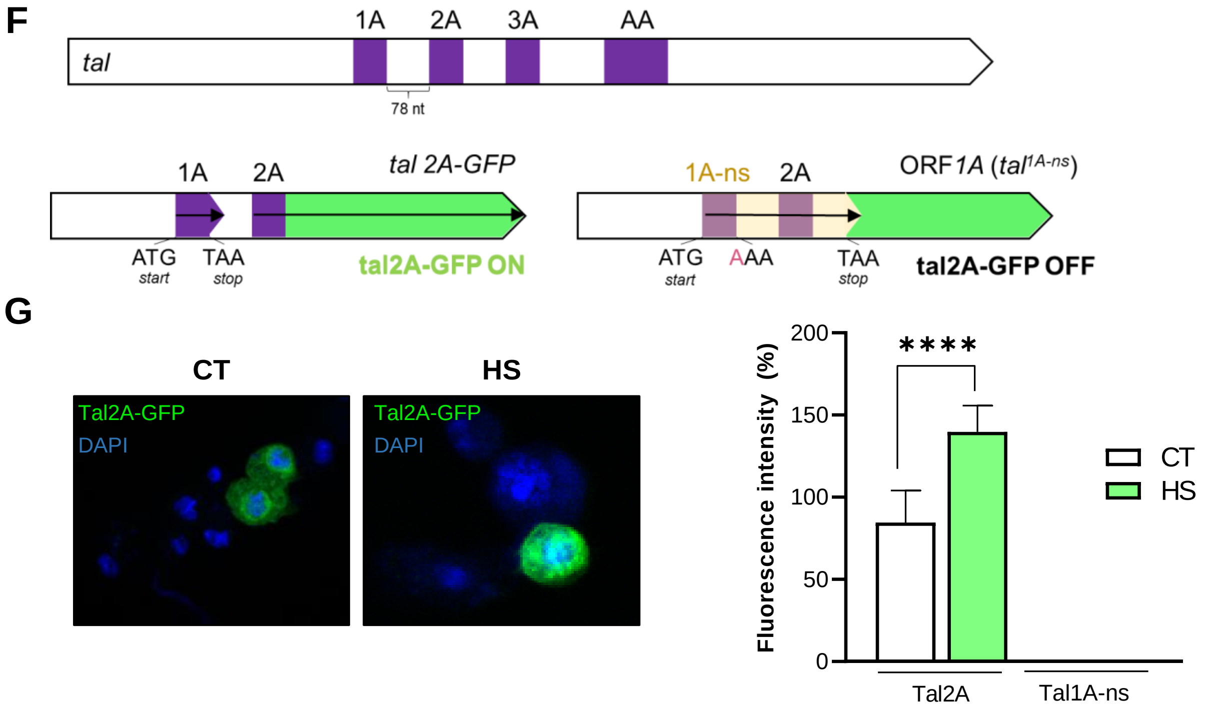
