## Supplementary figures for "Translation of small ORFs in non-canonical RNAs provides resilience to heat shock stress": Fig_S6.pdf

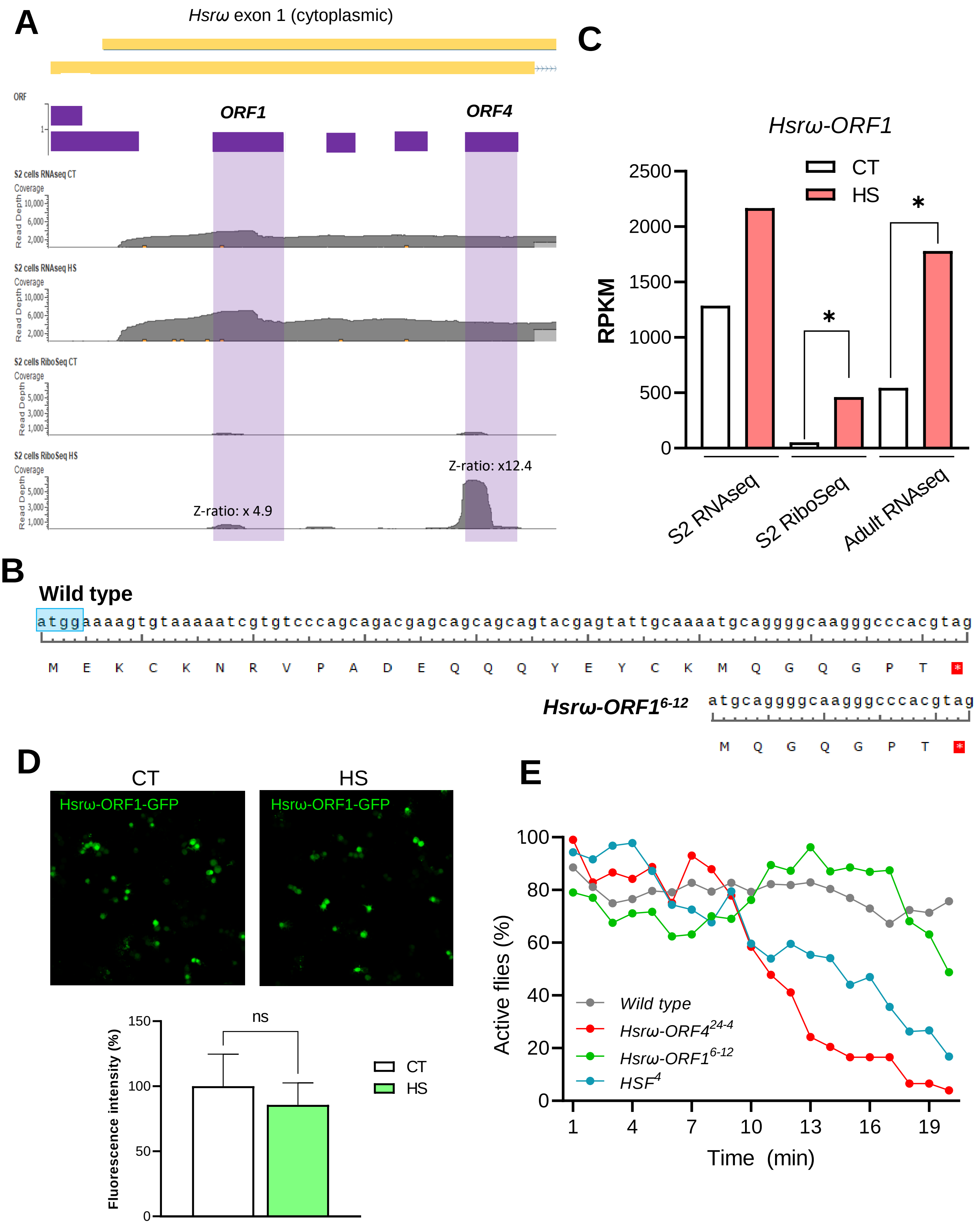


# F

| | HSF⁴/CyO; Hsrω-ORF4²⁴⁻⁴ x HSF⁴/CyO; Hsrω-ORF4²⁴⁻⁴ | HSF⁴/CyO x HSF⁴/CyO | HSF⁴/CyO; Hsrω-ORF1⁶⁻¹² x HSF⁴/CyO; Hsrω-ORF1⁶⁻¹² |
|---|---|---|---|
| *$HSF^4/CyO$* | 43 (100%) | 42 (100%) | 89 (100%) |
| *$HSF^4/HSF^4$* (expect ½ of CyO) | 4 (20%)* | 20 (100%) | 50 (100%) |
