## Supplementary figures and images for "Translation of small ORFs in non-canonical RNAs provides resilience to heat shock stress"

### Fig_S1.pdf

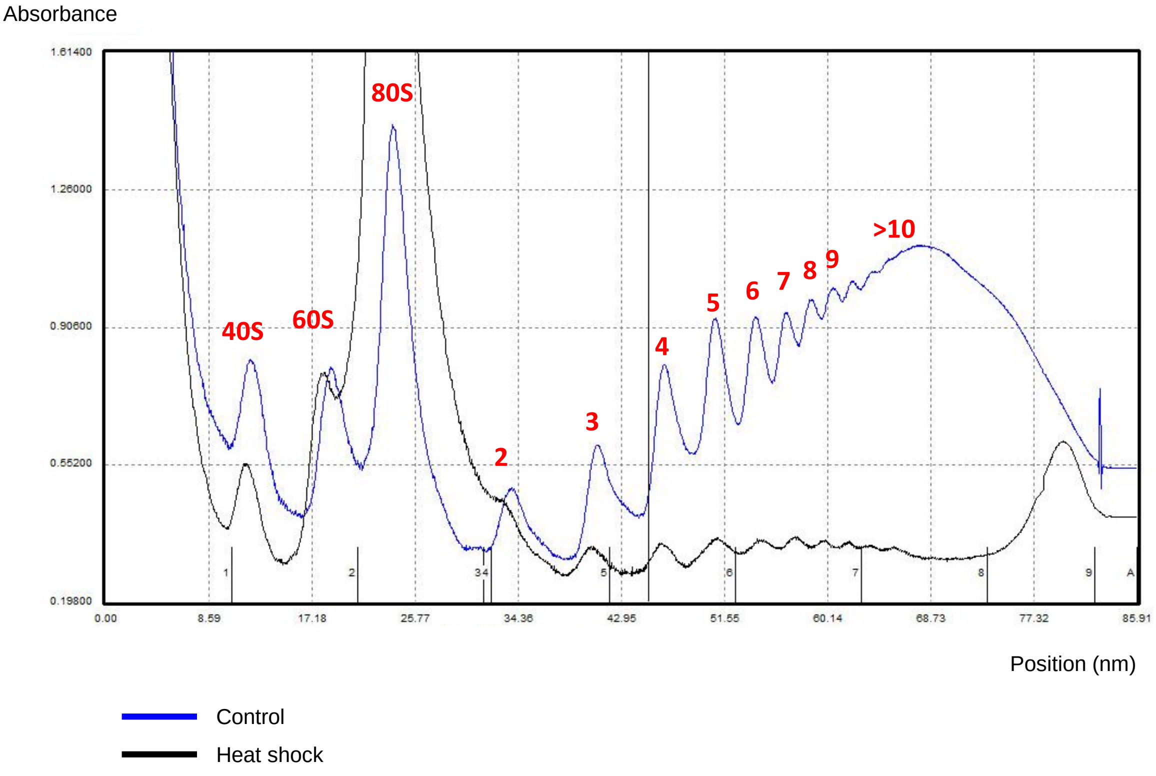
Absorbance
1.61400
1.26000
0.90600
0.55200
0.19800
0.00
8.59
17.18
25.77
34.36
42.95
51.55
60.14
68.73
77.32
85.91
40S
60S
80S
2
3
4
5
6
7
8
9
>10
Position (nm)
Control
Heat shock

### Fig_S3.pdf

**A**

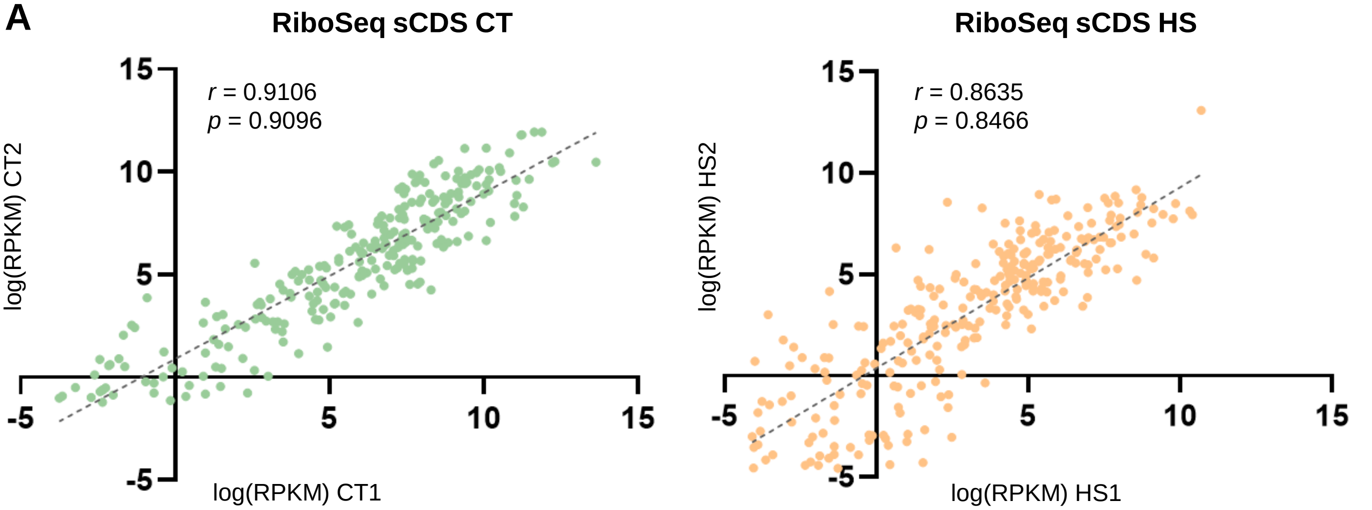


**B**

| sCDS counting | Only transcribed | Ribo-bound no translated | Translated |
|---|---|---|---|
| Control | 34 | 21 | 202 |
| Heat shock | 51 | 3 | 204 |

**C**

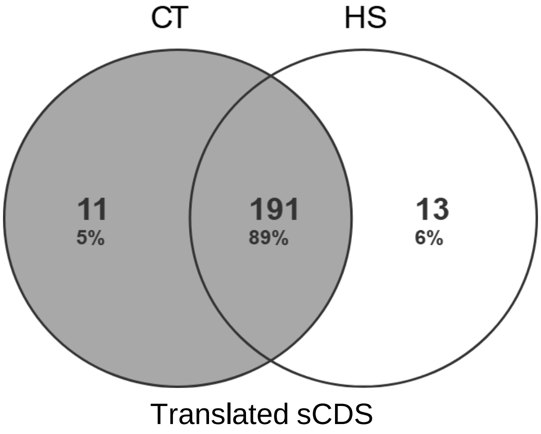


**D**

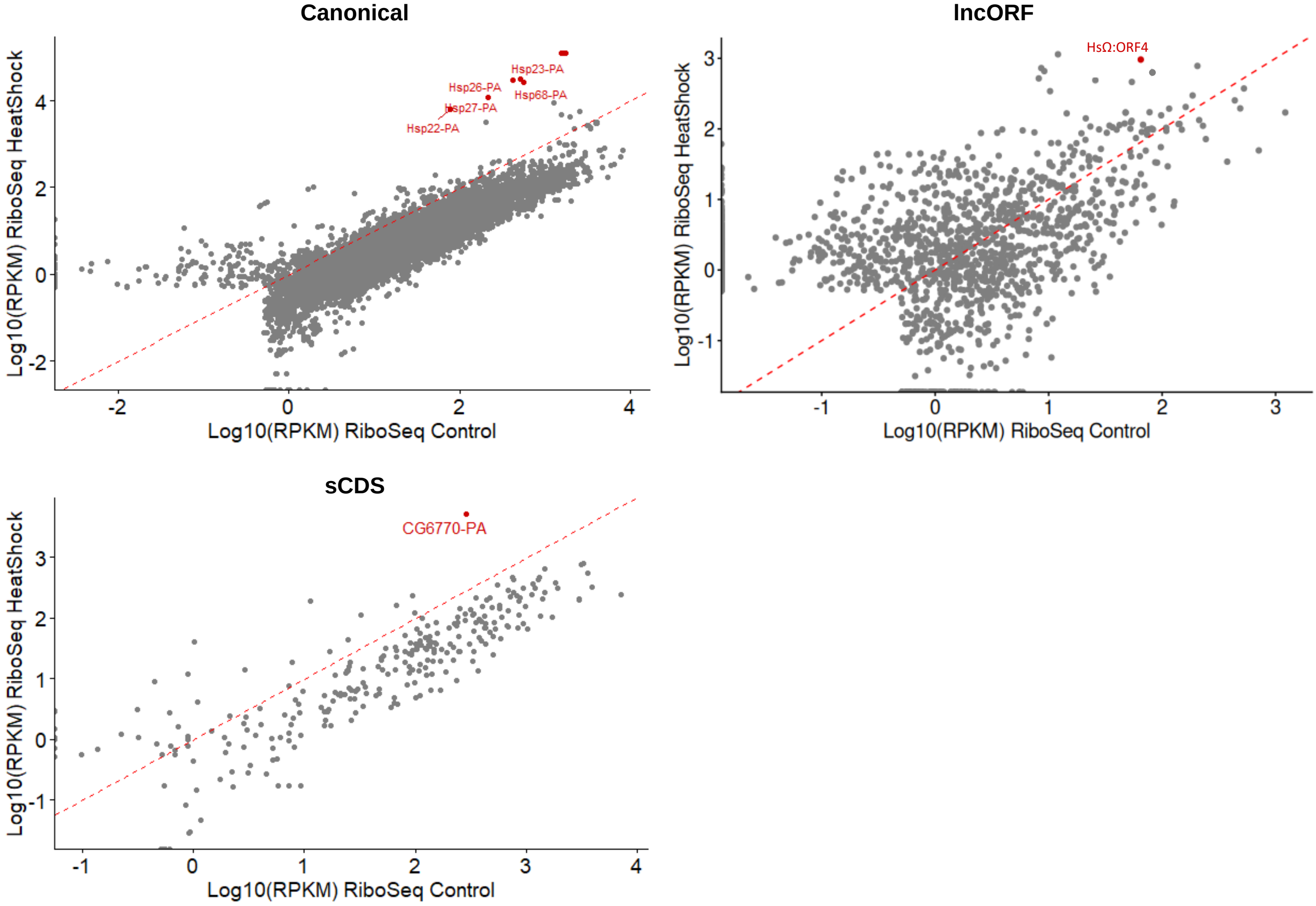

### Fig_S5.pdf

**A** **B**

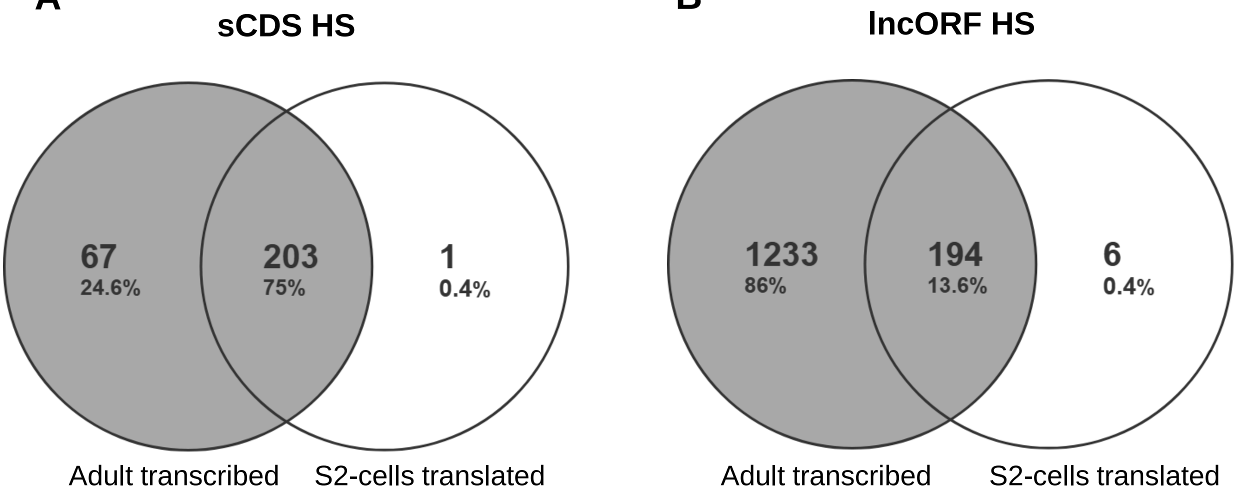


**C**

| ORF class | Up | | Down | | No significant changes (Z-ratio -1.5 < x <1.5) | | |
|---|---|---|---|---|---|---|---|
| **Canonical** | 11 | 0.1% | 284 | 2.3% | 11,972 | 97.6% | RPKM$^{FP}$ |
| **lncORF** | 12 | 0.7% | 18 | 1.1% | 1,668 | 98.2% | |
| **sCDS** | 1 | 0.4% | 7 | 2.5% | 269 | 97.1% | |

### Fig_S7.pdf

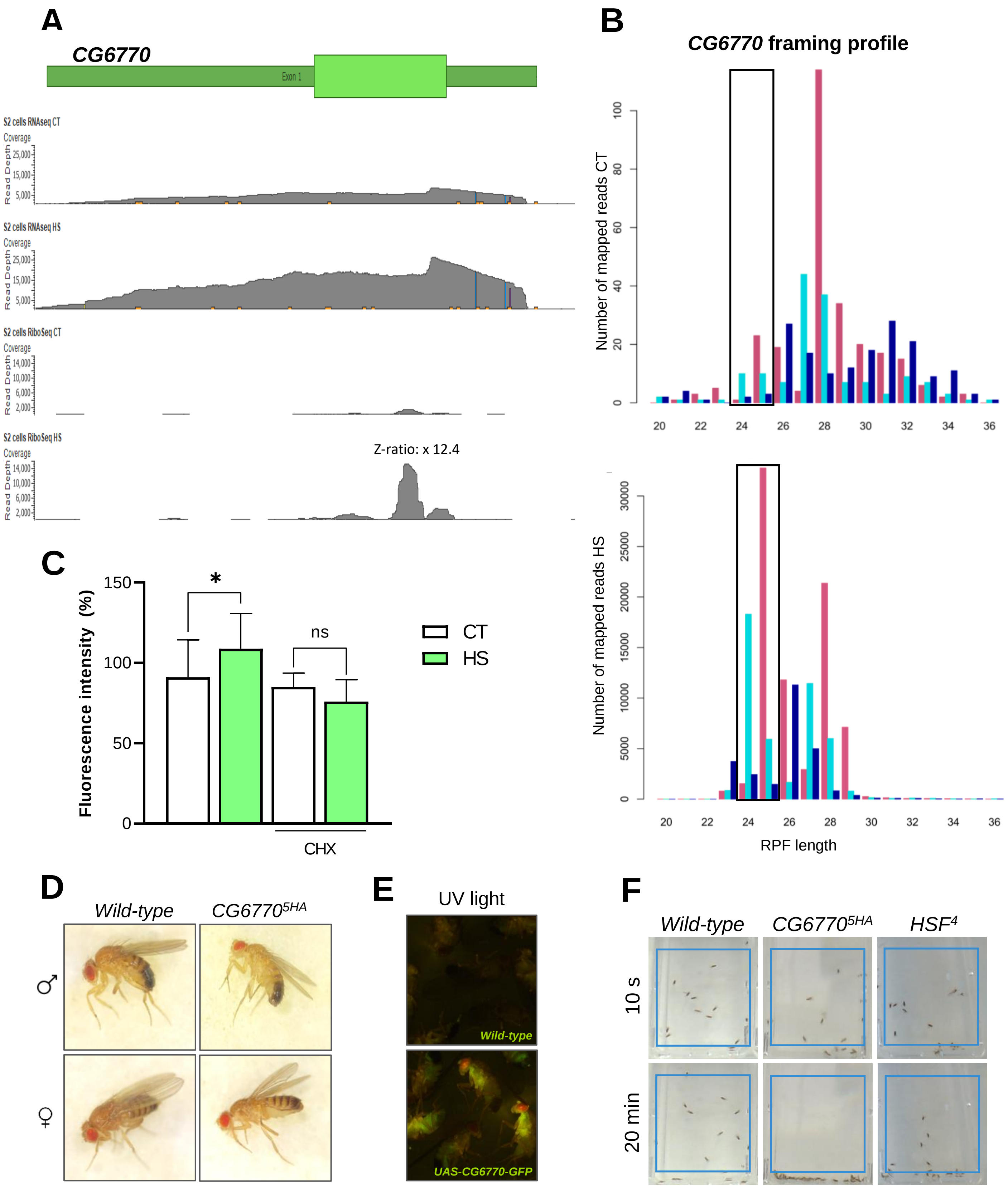


**G**

| | *CG6770*$^{5HA}$ x *HSF*$^{4}$/+ |
|---|---|
| *CG6770*$^{5HA}$ /+ | 132 (100%) |
| *CG6770*$^{5HA}$/*HSF*$^{4}$ | 0 (0%)* |
